# Novel transplantable mouse cell line model recapitulates invasive lobular breast carcinoma (ILC) phenotype and immune microenvironment

**DOI:** 10.64898/2026.07.30.741815

**Authors:** Sayali Onkar, Daisong Liu, Darcie Seachrist, Jian Zou, Christopher Merkel, Insa Thale, Alexander Chih-Chieh Chang, Linda Klei, Jian Chen, Kristen Weber Bonk, Kai Ding, Laura Savariau, Megan Yates, Jagmohan Hooda, Laura Stabile, Laura Rigatti, Peter C. Lucas, George Tseng, Keri Ruth, Creg J Workman, Adrian V. Lee, Dario A.A. Vignali, Steffi Oesterreich

## Abstract

Invasive lobular breast carcinoma (ILC) is the most common special histological subtype of breast cancer, which accounts for 10-15% of all cases. To study the phenotype characteristics, metastatic growth kinetic and immune microenvironment of ILC, we developed an orthotopically transplantable cell line model from the spontaneous mammary fat pad tumor of *CDH1-PTEN* dual knockout C57BL/6 mouse with Cre-loxP system, designated CPT6. CPT6 recapitulates single-file growth pattern of human ILC, with pleomorphic features and a high mitotic index. RNA sequencing together with whole exome sequencing reveals a luminal A subtype with targetable driver mutations such as Kras G12C. As a novel orthotopically transplantable ILC model in immune competent mice, CPT6 shows robust *in vivo* growth and metastatic rate, and has moderate immunogenicity which appears to be T-cell independent. We also profiled the immune microenvironment of CPT6, revealing a myeloid-rich environment with dominant M2-macrophage population, which is concordant with human ILC. In summary, this model recapitulates human ILC phenotype and represents a valuable preclinical platform for evaluating immunotherapy and other therapeutic strategies for invasive lobular breast carcinoma.

## Introduction

Invasive lobular breast cancer (ILC) is the second most common histological subtype of breast cancer that accounts for 10-15% of all breast cancer ^[1]^. It is characterized by a single-file growth pattern resulting from loss of E-cadherin (*CDH1*), which mediates cell-cell adhesion. Thus, ILC typically exhibits a diffuse and infiltrative growth pattern. Clinically, because 90% of ILC are hormone receptor positive (HR+) ^[2]^, patients with ILC are mostly treated with endocrine therapy and CDK4/6 inhibitors, and show worse long-term survival as compared to the most common subtype invasive ductal carcinoma/carcinoma of no special type (IDC/NST) ^[3]^. On a molecular level, ILCs are commonly estrogen receptor positive (ER+) and exhibit predominantly luminal A PAM50 subtype, with enriched mutations in *PI3K, PTEN*, *TBX3*, *FOXA1* etc as compared to NST ^[4]^, indicating distinct origins and genomic drivers in this subtype. Within ILC, 10-15% are identified as invasive pleomorphic lobular carcinoma (pILC) characterized by nuclear pleomorphism, increased nuclear dimension, intense mitotic activity and vacuolation ^[5]^. pILC exhibits aberrant genomic alterations including chromosomal copy number variation and structure variations ^[6]^ and also shows high growth rate and worse prognosis clinically ^[7]^. Besides intrinsic features of ILC, extrinsic factors, especially the tumor microenvironment, also play a vital role in the progression and metastasis of ILC. In non-immune microenvironment, ILC shows distinct cancer associated fibroblasts (CAFs) markers expression compared with NST, like FAP-α and FSP-1/S100A4 which are associated with poor prognosis ^[8]^. In the immune microenvironment, ILC shows a macrophage-dominant immune cell infiltration, with a predominant M2-like protumor macrophage constitution when compared to NST ^[9]^. Thus, targeting the macrophage-dependent immune microenvironment of ILC as combination strategy might a promising approach in treatment of ILC. Therefore, in order to study *in vivo* characteristics of ILC, it becomes vital to develop a robust syngeneic animal model that recapitulates human ILC intrinsic features as well as its tumor microenvironment.

Currently, three genetically engineered mouse models (GEMM) of ILC have been described, all involving deletion of *CDH1*, combined with either co-deletions of *TP53,* co-deletion of *PTEN*, or an activating mutation of *PIK3CA* ^[10][11][12]^. Mammary-specific inactivation of *TP53* and *CDH1* leads to the formation of ER negative pleomorphic invasive lobular carcinoma ^[10]^. Cells derived from this spontaneous tumor also showed tumorigenicity when orthotopically transplanted in immune incompetent mice. Concurrent inactivation of *CDH1* and activation *PIK3CA* leads to formation of ER/PR positive classical ILC tumor *in vivo* ^[11]^, and finally co-loss of *CDH1* and *PTEN* in FVB/N mouse background caused formation of spontaneous tumors that recapitulate histological feature of classical ILC with ER expression and that are responsive to PI3K inhibition ^[12]^. However, no stable and transplantable GEMM derived cell models have been developed for immune competent mice to enable testing of immunotherapy and other targeted therapy.

In this study, we developed and characterized an ILC cell line from a mammary tumor arising after dual deletion of *CDH1* and *PTEN* through Cre-loxP system in a C57BL/6 background. The line was designated CPT6, an acronym for <u>C</u>DH1-<u>P</u>TEN-<u>T</u>UMOR-Bl<u>6</u>. We profiled the phenotypic, genomic and transcriptomic features of CPT6, and investigated primary and metastatic growth, estrogen response, as well as the immune microenvironment of this model. CPT6 is a novel luminal A ILC model that is orthotopically transplantable into immune competent mice. This model preserves the molecular and histological features as human ILC and recapitulates composition of tumor microenvironment in human ILC. The CPT6 model thus provides a valuable platform for future preclinical testing of immunotherapy and other combination treatment strategies for invasive lobular breast carcinoma.

### CPT6 Cell Line Exhibits ILC Histological and Molecular Features

C57BL/6 is one of the most widely used inbred mouse strains with known genotypic features and established biological tools. Therefore, we selected C57BL/6 as the genetic background and bred *PTEN*^L/L^ x *CDH1*^L/L^ x *WAP*^Cre^ mice to achieve the desired genetic modification (**Figure 1a)**. Notably, WapCre males were crossed with wild-type (WT) females, as WapCre females fail to produce viable litters, resulting in neonatal mortality. A spontaneous mammary tumor was isolated from a 32-week-old female mouse; tumor tissue was enzymatically digested and EpCAM-positive (EpCAM⁺) cells were sorted to establish the CPT6 cell line (**Figure 1b**). Flow cytometry revealed 60.7% EpCAM +ve cells, which were recognized as epithelial cells and subsequently sorted (**Figure S1a**). The sorted cell line retained tumorigenic capacity following orthotopic mammary fat pad injection into immunocompetent C57BL/6 mice, enabling further investigation of *in vivo* growth dynamics and immune profiling.

**Figure 1.**
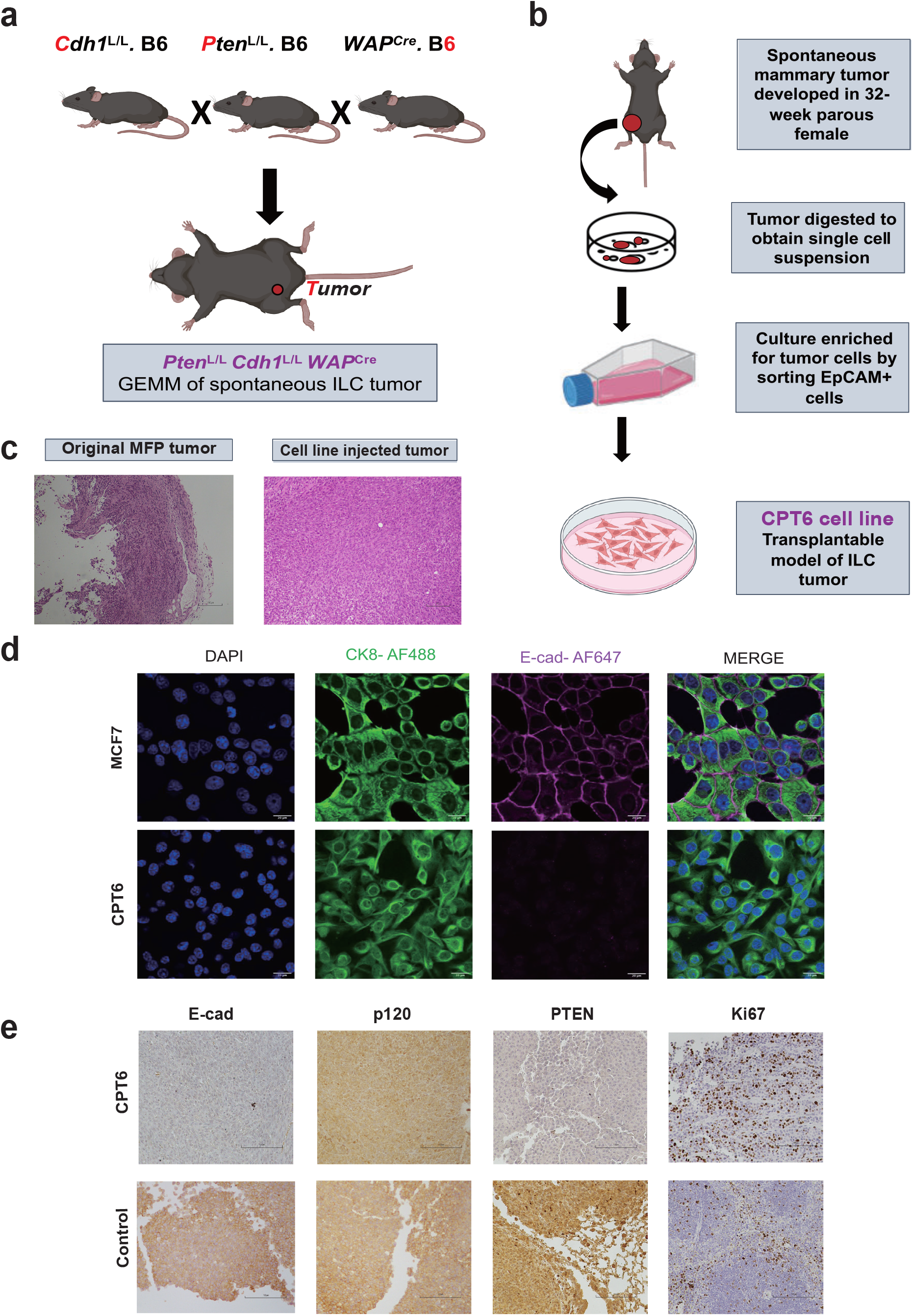
Generation and cellular characterization of CPT6 cell line**. a.** Breeding scheme for generating PtenL/L Cdh1L/L WapCre mouse model of spontaneous breast lobular tumor. GEMM: Genetically engineered mouse model. **b.** Schematic representation of CPT6 cell line generation from mouse mammary tumor. **c.** H&E staining (20x) of spontaneous tumor (left) and CPT6 cell line injected tumor (right). Cancer cells shows evident single file growth pattern confirmed by pathologists. **d.** Immunofluorescence images (40x) of MCF7 (human, E-cad +ve, top row) and CPT6 cell line (bottom row) with luminal cytokeratin CK8 (green) and E-cadherin (purple) in dual staining. **e.** Immunohistochemistry images (20x) of CPT6 and control cell pellets. SSM3 cell pellets were used for control of E-cad and p120 staining. PTEN +ve control used mouse lung and Ki67 +ve control used mouse lymph node.

To confirm that the spontaneous tumor had acquired the phenotypic and molecular features of classical ILC, we performed H&E staining of both the original spontaneous tumor and the tumors generated by orthotopic injection. The cells in both cases displayed the typical "single file" growth pattern characteristic of ILC, confirmed by two pathologists (**Figure 1c**). The *in vivo* tumor from injection also showed features of densely packed malignant cells with high mitotic rates, and some spindle as well as moderate pleomorphic morphology. These findings establish CPT6 as a syngeneic, orthotopically transplantable mouse ILC cell line with pleomorphic features suitable for use in immunocompetent hosts.

To characterize the *in vitro* features of CPT6, we first performed immunofluorescence (IF) staining of CPT6 compared to the human MCF7 cell line (E-cadherin +ve), using E-cadherin and cytokeratin 8 (CK8) as biomarkers (**Figure 1d**). Results showed positive CK8 staining in both CPT6 and MCF7, indicating a luminal origin for these two cell lines. As expected, MCF7 but not CPT6 displayed robust positive E-cadherin staining at the cell membrane.

To further characterize the molecular features of CPT6, we performed immunohistochemistry (IHC) staining of E-cadherin, p120, PTEN and Ki67 (**Figure 1e**). Strong membranous E-cadherin staining was observed in the mouse SSM3 cell line (NST subtype), whereas no staining was detected in CPT6. As expected, p120 catenin, which normally associates with E-cadherin at the cell membrane and loses its membrane anchor upon E-cadherin loss, displayed strong membranous staining in SSM3 but diffuse cytoplasmic redistribution in CPT6. These IHC findings indicate the absence of functional E-cadherin in CPT6 and the cytoplasmic re-localization of p120, which was corroborated by Western blot analysis confirming complete loss of E-cadherin protein in CPT6. As a control for mouse E-cadherin, we used parental SSM3 cells and CRISPR-Cas9-mediated CDH1-knockout SSM3 cells (**Figure S1b**). PTEN loss in CPT6 was similarly confirmed by both IHC (**Figure 1e**) and Western blot analysis (**Figure S1b**), which showed absent staining relative to positive controls. CPT6 cells displayed high proliferative activity as reflected by a Ki67 positivity of 40.6% through QuPath analysis (**Figure 1e)**. This was further supported by cell cycle analysis using propidium iodide (PI) staining, which revealed a high S-phase fraction of 25.7% (**Figure S1c**).

### Genomic Characterization Establishes CPT6 Similarity to Human ILC Cell Lines with Targetable Driver Mutations

Next we characterized the transcriptome of CPT6 by performing bulk RNA sequencing with early passage CPT6 cells grown *in vitro*. For comparison, we used the mouse E0771 cell line, also derived from tumors from C57BL/6 mice. There are different reports on ER expression levels in E0771 ^[13][14]^, and it has been considered to be of luminal B in PAM50 subtype analysis ^[15]^ but has also been described to be more TNBC-like ^[16]^.

We first combined the transcriptomes of CPT6 and E0771 to determine PAM50 subtype classification. We used RNA sequencing data from human ILC cell line encyclopedia (ICLE) with PAM50 subtype as reference ^[17]^. We observed that CPT6 transcriptomes clustered closer to the luminal A subtype compared to E0771 (**Figure 2a**). We also sequenced *in vivo* tumors formed by CPT6 and E0771 mammary fat pad injection to assess how the tumor microenvironment might influence the transcriptome. We compared the two mouse tumor transcriptomes with human reference from combined datasets, and t-SNE clustering confirmed that CPT6 aligned more closely with the LumA subtype, while E0771 clustered nearer to the LumB subtype (**Figure 2b**).

**Figure 2.**
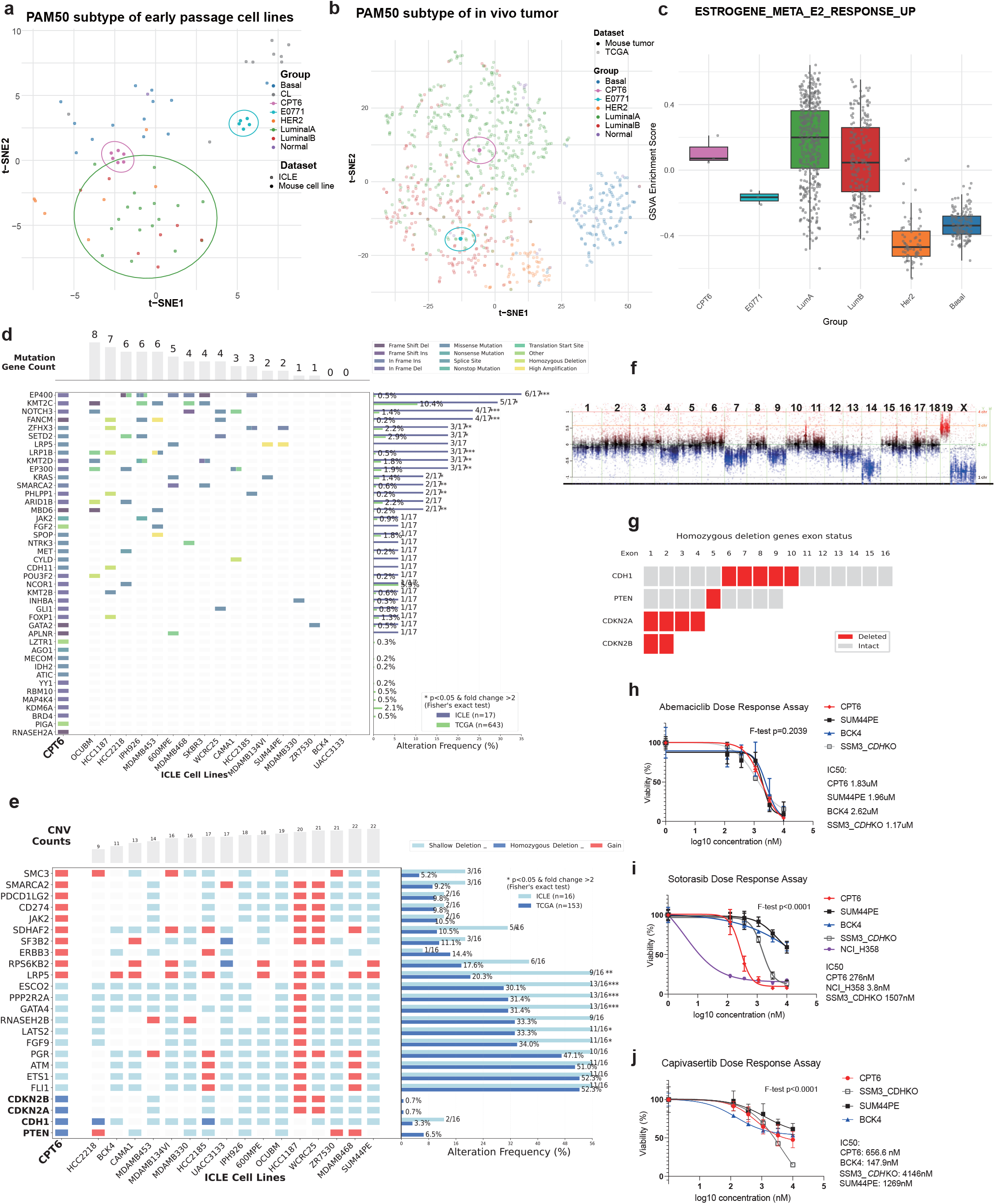
Molecular characterization of CPT6 transcriptome and genome**. a.** t-SNE clustering of PAM50 subtype in early CPT6 passage in comparison to E0771, ICLE and in house mouse dataset. **b.** Pathway enrichment score comparison of Estrogene_Meta_E2_Response_Up pathway in CPT6 and E0771 compared to TCGA data with denoted PAM50 subtypes (Basal, HER2, LuminalA, LuminalB). **c.** t-SNE clustering of in vivo CPT6 and E0771 PAM50 subtype in comparison to TCGA dataset. **d.** Oncogenic mutation profile of CPT6, comparing with mutation type of ICLE (human ILC cell lines) and mutation frequency of ICLE and TCGA ILC samples. **e.** Top 20 CNV profile of CPT6, comparing with CNV type of ICLE and CNV frequency of ICLE and TCGA ILC samples. **f.** Chromosomal copy number variation of CPT6. **g.** Exon status of homozygously deleted genes in CPT6 genome. **h.** Dose response assay to Kras G12C inhibitor sotorasib. SSM3 CDH KO is achieved via CRISPR-Cas9 knockout. NCI_H358 is a non-small cell lung cancer cell line with known Kras G12C mutation. **i.** Dose response assay to CDK4/6 inhibitor abemaciclib. **j.** Dose response assay to Akt inhibitor capivasertib.

Given the LumA subtype, we next measured ER expression and signaling activity in CPT6 cells. Although western blot analysis revealed negligible levels of ERα expression (**Figure S2a**), ER signaling pathway was active, as reflected in an enrichment of the E2 response signature ^[18]^ similar to LumA/LumB tumors (**Figure 2c**). This was supported by analysis of *ESR1* mRNA expression, that was closer to LumA/LumB tumor compared to that of E0771 (**Figure S2b**). Functional assays for ER transcriptional activity showed minimal responses, limited to induction of Greb1 and PR (**Figure S2c**), but without effects in reporter assays (**Figure S2d**) or *in vitro* (**Figure S2e-f**) and *in vivo* (**Figure S2g-h**) growth. We also attempted to increase estrogen response through overexpression of ER, but initially successful overexpression of ER was lost with increasing passaging in culture (**Figure S2i**), and the overexpression did not mediate estrogen response in growth (Figure **S2j**). Collectively, these results demonstrate that CPT6 cells and *in vivo* tumors preserve luminal features, with CPT6 showing closer LumA resemblance characterized by an overall active estrogen response pathway activity, despite limited estrogen response as measured by growth.

To characterize the genomic alterations of CPT6, we performed whole exome sequencing (WES), using liver cells from healthy C57BL/6 mice as controls. We first profiled high-impact oncogenic mutations with human gene homologs and compared the mutation profile with ICLE ^[17]^ and ILC samples in The Cancer Genome Atlas Program (TCGA) ^[19]^ (**Figure 2d**). We identified mutations in 42 genes, with 30 appearing in ICLE and 32 appearing in TCGA ILC tumors. Among these mutated genes is KTM2C, that has high mutation rates in both TCGA (7%) and METABRIC (14%). As expected from a cell line, mutation rates of genes mutated in CPT6 were higher in ICLE compared to TCGA for genes co-occurring in both datasets. This demonstrates that the oncogene profile in CPT6 resembles human ILC cell lines and tumors. Notably, we analyzed the pathogenicity of all somatic mutations and variant allele frequency and identified heterozygous *Kras* G12C and homozygous *Cdh11* E462D as driver mutations for CPT6 oncogenesis (**Figure S3a**).

We next profiled all somatic mutations (Supplementary Material 03), and by analyzing the variant allele frequency together with copy number variations, we observed that CPT6 somatic mutations clustered into three subclones by genomic composition (**Figure S3b**). Reactome analysis showed an enrichment of somatic mutations in processes involving metabolism, signal transduction, disease, and protein modification (**Figure S3c).** The overall TMB in CPT6 was similar to other cell lines in ICLE and significantly higher than TCGA (**Figure S3d**).

Analysis of most frequent copy number variation (CNV) showed gains and deletions shared with both ILC cell lines in ICLE, and tumors in TCGA (**Figure 2e**). Profiling of chromosomal CNV pattern revealed aberrant chromosomal changes, featuring whole-chromosome shallow deletions of chr7, 14, and X, as well as whole-chromosome amplification of chr19 (**Figure 2f**). *CDH1*, *PTEN*, *CDKN2A* and *CDKN2B* showed homozygous deletion (**Figures 2e-g**). *CDH1* exhibited deletion of exons 6-10, while *PTEN* showed exon 5 deletion, confirming expected *CDH1* and *PTEN* deletions resulting from Cre-loxP gene targeting approach. As the first exons remained intact, some RNA can be transcribed, but no functional protein is present (**Figures 1e and S1b**).

*CDKN2A* (p16INK4a/p19ARF) and *CDKN2B* (p15INK4b) are located in close proximity on Chromosome 4. There is complete loss of this region, including all exons, potentially pointing towards a role in CPT6 tumorigenesis. Given the known role of *CDKN2A/B* as inhibitors of CDK4/6, we asked whether CPT6 cells might show enhanced sensitivity to CDK4/6 inhibitor abemaciclib. As controls, we included two human ILC cell lines: SUM44PE (*CDKN2A/B* shallow deletion) and BCK4 (*CDKN2A/B* diploid), one mouse cell line SSM3 with CRISPR-mediated *CDH1* knockout (*CDKN2A/B* diploid). There was no significant difference in the IC50 between the cell lines (**Figure 2h**) suggesting that loss of CDKN2A/B does not affect sensitivity to CDK4/6 inhibitors in this model.

We also tested whether the KrasG12C mutation would affect response to sotorasib. In addition to KRAS WT ILC/ILC-like cell lines (SUM44PE, BCK4, and SSM3_CDHKO), we included a human lung adenocarcinoma cell line NCI-H358 that has the same KrasG12C mutation as CPT6. We observed significantly higher sensitivity in CPT6 compared to the KRAS WT cell lines although they were not as sensitive as the KrasG12C-mutated lung cancer cells (**Figure 2i**).

Since CPT6 has a PTEN deletion, we tested its response to PI3K/Akt pathway inhibition using the AKT inhibitor capivasertib. The inhibition in CPT6 was stronger compared to SUM44PE and SSM3_CDHKO that do not have PI3K/Akt pathway alteration, but weaker compared to BCK4 that harbors a *PIK3CA* H1047R mutation (**Figure 2j**).

### CPT6 displays rapid *in vivo* growth and metastases to the lungs

Orthotopic injection of CPT6 cells results in rapid growth similar to that of E0771 cells (**Figure 3a**). For both cell lines, tumor growth was initially slow until day 15 post-injection, after which tumors grew exponentially and reached terminal size between days 28-35 post injection. Notably, both tumor types presented varying degrees of ulceration with onset between days 18 and 21. In some cases, mice required euthanasia due to ulceration before reaching terminal tumor size, which occurred more frequently in the CPT6 cohort compared to the E0771 cohort.

**Figure 3.**
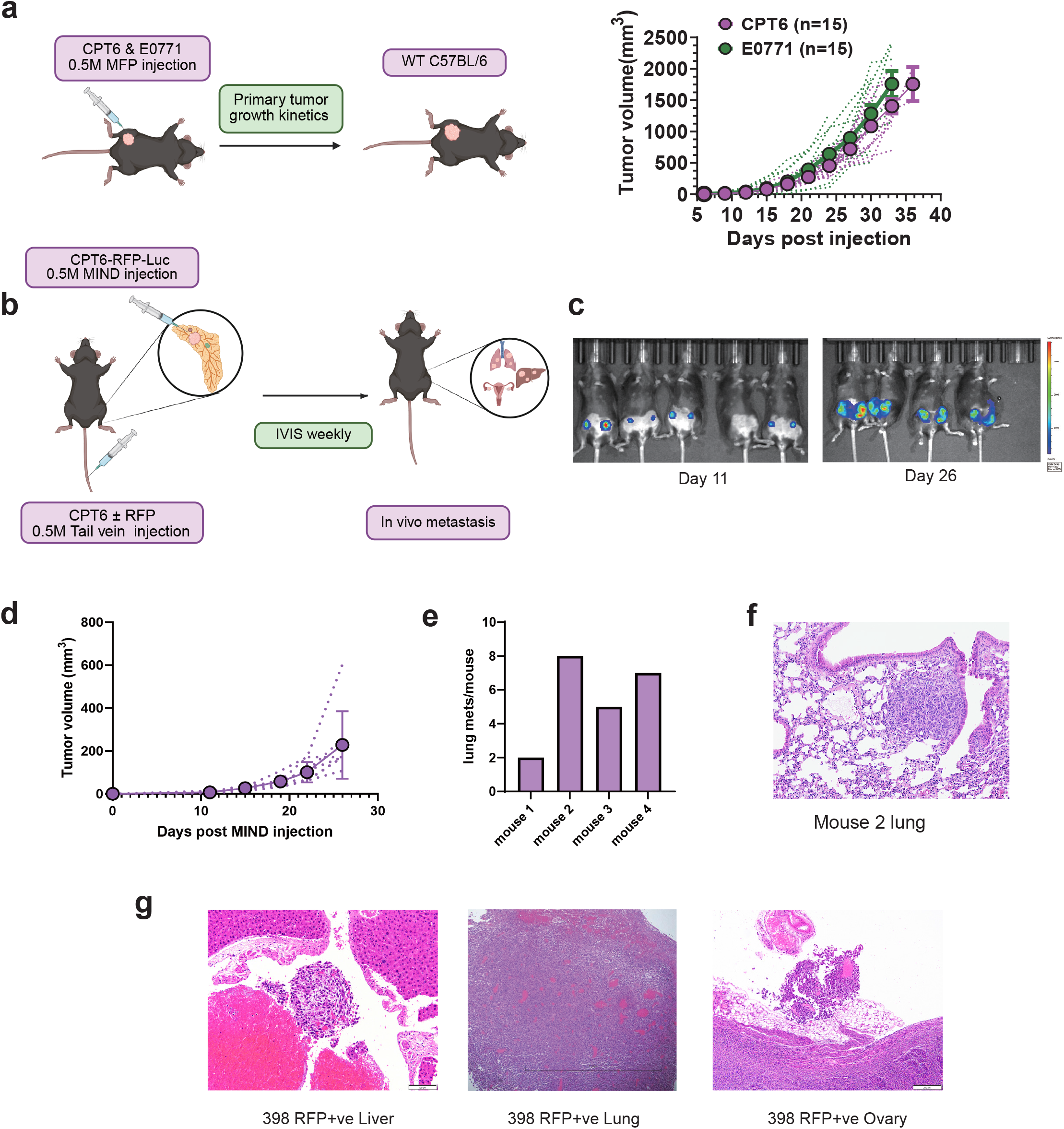
In vivo growth and metastatic kinetic of CPT6**. a.** Scheme of in vivo CPT6 & E0771 MFP injection (left) and in vivo growth rate comparison of CPT6 and E0771 in WT C57BL/6 (right) **b.** Scheme of MIND (mouse intraductal) and tail vein injection of CPT6 cells in vivo. **c.** IVIS results of primary tumor growth size post MIND injection on Day 11 (left) and Day 26 (right). **d.** Primary tumor volume growth dynamic post MIND injection. **e.** Number of lung micrometastasis post MIND injection in H&E section at time of sacrifice (n=4). **f.** H&E staining of metastatic sites post MIND injection lung (20x). g. H&E staining (4x) of metastatic sites (liver, lung and ovary) in RFP-ve CPT6 cell tail vein injection in mouse 398.

Rapid growth rate in the mammary fat pad prevented us from determining potential metastases after MFP injection. We therefore performed mammary intraductal (MIND) and tail vein (TV) injection in C57BL/6 mice using red fluorescent protein (RFP)-labeled CPT6 cells and conducted weekly monitoring via In Vivo Imaging System (IVIS) (**Figure 3b**). Overall, CPT6 demonstrated rapid growth, which was readily visualized by IVIS imaging (**Figure 3c**). The growth kinetics of primary tumors following CPT6 MIND injection were similar to those observed with direct mammary fat pad injection, as shown in **Figure 3d**. Microscopic analysis of H&E staining of lung section at day 28 revealed lung metastases in four out of five mice (**Figure 3e**), with an average of four metastatic foci per section per mouse. A representative IHC of a CPT6 lung metastasis is shown in **Figure 3f**.

For the TV injections, where we injected both unlabeled and RFP-labeled CPT6 cells, lung metastases was observed in 100% of the injected mice between days 14-21, regardless of RFP label status. In one of the mice (#398) injected with RFP-labeled cells, metastasis to the ovary and liver were observed as well (**Figure 3g**). Overall, CPT cells have a robust ability to metastasize to the lung, with less frequent metastases to other sites including liver and ovary.

### CPT6 demonstrates Intermediate Immunogenicity, characterized by myeloid cell infiltration

A main goal for the generation of a syngeneic ILC model was to generate a transplantable model that will help the ILC research community to understand interaction with the tumor microenvironment (TME) especially the immune system. As a first step, we analyzed MHC class I surface expression on the cell line as a surrogate marker for tumor cell immunogenicity (**Figure 4a**). We included B16 melanoma cells (poor immunogenicity) and MC38 colon carcinoma cells (high immunogenicity) alongside E0771 as controls ^[20]^. CPT6 cells displayed minimal to absent H2Db expression and moderate H2Kb expression. In contrast, E0771 cells expressed higher levels of both H2Db and H2Kb haplotypes. We concluded that CPT6 cells possess intermediate immunogenicity, greater than B16 but lower than E0771 or MC38.

**Figure 4.**
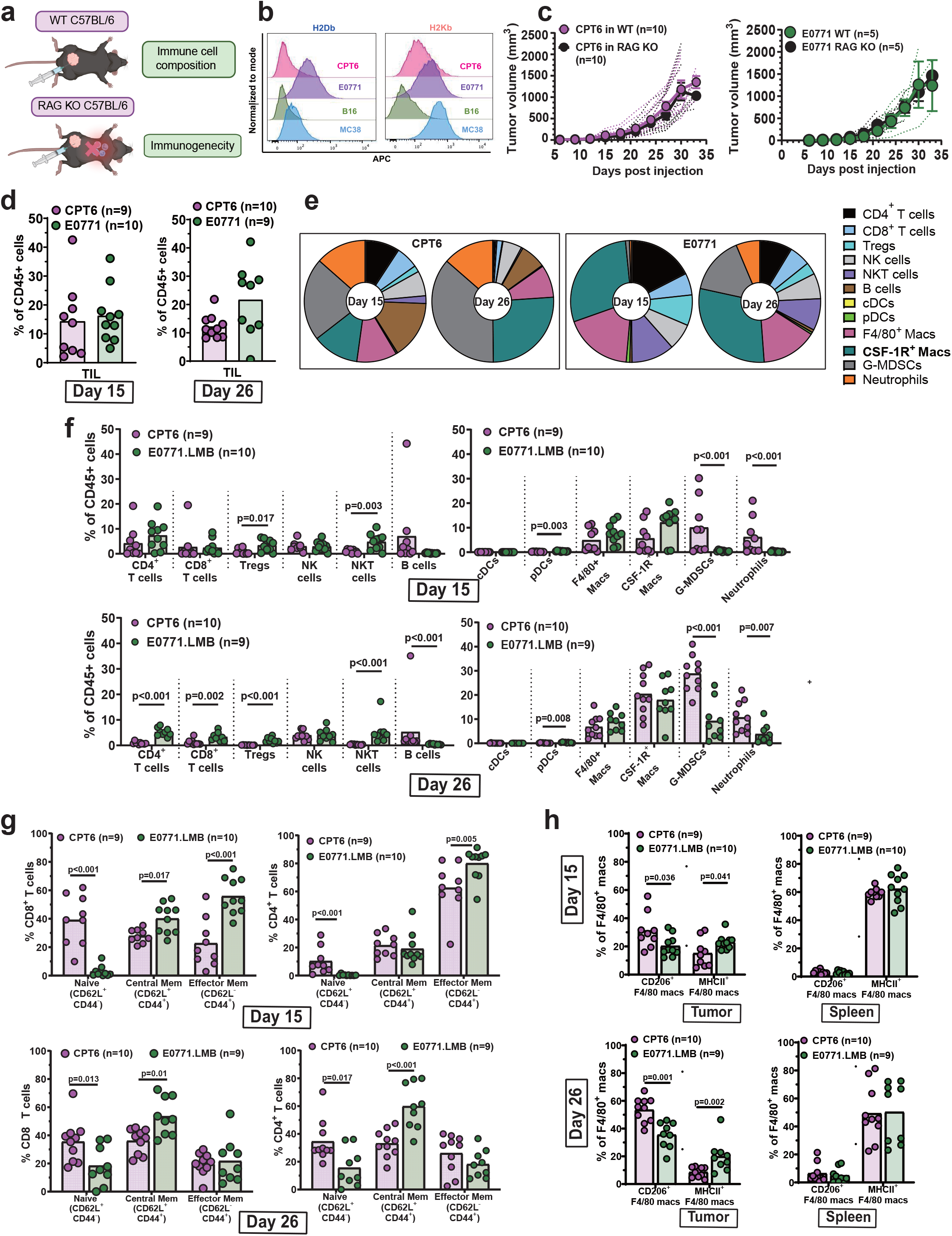
Immune infiltration landscape of tumor microenvironment in CPT6 and E0771 in vivo tumor. **a.** Scheme of in vivo CPT6 & E0771 MFP injection in WT C57BL/6 and RAG KO mice. **b.** Histogram comparing cell surface expression of H2Db (left) and K2Kb (right) MHC class I haplotypes on CPT6, E0771, B16 and MC38 cell lines with flow cytometry. **c.** In vivo growth rate comparison of CPT6 (left) and E0771 (right) in WT and RAG KO C57BL/6. **d.** Tumor infiltrating lymphocyte (TIL) proportion in CD45+ cells at Day 15 (left) and Day 26 (right) in CPT6 and E0771 in vivo tumor. **e.** Pie chart comparing total tumor infiltrating immune cells composition of CPT6 at Day 15 (left) and Day 26 (right). **f.** Comparison of lymphoid (left) and myeloid (right) immune cell infiltration between CPT6 and E0771 at Day 15 (upper panel) and Day 26 (lower panel). **g.** Frequency of naive, central memory and effector memory phenotypes of tumor infiltrating CD8+ and CD4+ T cells in CPT6 and E0771 at Day 15 (upper) and Day 26 (lower). **h.** Comparison of macrophage phenotypic subtypes: CD206+ (M2-like macrophage) and MHCII+ (M1-like macrophage) in CPT6 and E0771 tumors and spleens at Day 17 (up) and Day 26 (down).

To determine the effect of the immune microenvironment on CPT6 growth, we injected CPT6 and E0771 cells into the mammary fat pads of wild-type (WT) and RAG knockout (KO) C57BL/6 mice (**Figure 4b**). We observed similar growth kinetics for both CPT6 and E0771 in WT and RAG KO mice, suggesting that lymphocytes, namely T and B cells, may not play a critical role in the development of these mammary tumors (**Figure 4c**).

This stimulated us to comprehensively profile the TME, particularly with respect to the myeloid compositions. Since the model affords assessment of the temporal evolution of this immune response, we chose day 15 (D15) post injection as an early timepoint (representing low tumor burden, active immunoediting) and day 26 (D26) as a mid-late time point (representing high tumor burden, potential immune escape). At the early timepoint on D15 total immune infiltration (CD45^+^ cells) in CPT6 tumors was found to be comparable to that in E0771 tumors (**Figure 4d**, left panel). However, as the tumors progressed to D26 (**Figure 4d**, right panel), the median percentage of immune cells was unaltered in CPT6 (∼12-15%) compared to E0771 tumors, which demonstrated a small increase in the percentage of infiltration (∼15% at D15 to ∼22% at D26). We also compared the CD45+ cell proportion in the spleen and lymph node at D15 and D26, and the proportions are not significantly different in both timepoint at both locations (**Figure S4a**).

We next compared the proportions of lymphoid and myeloid cells at D15 and D26 (**Figure 4e**) using an approach and marker combinations detailed in **Figures S4b.** As expected, CPT6 tumors harbored lower proportion of all lymphocytes (T and B cells, NK cells) at D15 compared to myeloid cells (DCs, macrophages, MDSCs). This balance was tipped further in favor of myeloid cells making the CPT6 TME practically devoid of T cells by D26. In contrast to CPT6, the E0771 TME harbored much greater proportion of lymphocytes at both D15 and D26, but followed a similar trend of reduction in lymphocyte populations and expansion of myeloid cells by D26. We then directly compared the individual lymphoid and myeloid cell frequencies in the two models at D15 (**Figure 4f, upper panel**) and D26 (**Figure 4f, lower panel).** In both CPT6 and E0771, T cells were less than 10% of all CD45^+^ cells but were significantly higher in E0771 compared to CPT6 by D26. In contrast to T cells, CPT6 tumors harbored a greater frequency of B cells at both early and late timepoints (p<0.001) compared to E0771 tumors. Within the myeloid compartment, CPT6 TME was dominated by granulocytic myeloid derived suppressor cells (G-MDSCs) and neutrophils, which were significantly higher than E0771 at both D15 and D26. On the other hand, macrophages (F4/80^+^ and CSF-1R^+^) dominated the myeloid infiltrate in E0771 tumors. We also observed a significant increase in CSF-1R^+^ tissue- resident macrophage (p<0.001) and G-MDSC infiltration (p=0.001) in CPT6 as the tumors progressed from D15 to D26. Finally, both TMEs were infiltrated by a very small percentage (<2%) of conventional dendritic cells (cDCs) and plasmacytoid dendritic cells (pDCs), the latter being significantly higher in E0771 both early and late timepoints.

### CPT6 TME harbors naïve and potentially exhausted T cells early during tumor growth, and immunosuppressive macrophages

Even though the percentage of T cells infiltrating CPT6 tumors was consistently low across both timepoints analyzed, we investigated the functional phenotype of T cells with the aim of identifying markers and potential strategies to impact T cell function. We first assessed the distribution of T cells between naïve, central memory and effector phenotypes for CD8^+^ (**Figure 4g, left panels**) and CD4^+^ T cells (**Figure 4g, right panels**) infiltrating CPT6 and E0771 tumors at Day 15 (**Figure 4g, top panels)** and Day 26 (**Figure 4g, bottom panels)**. In the CD8 T cell compartment, 40% of CPT6 tumor infiltrating cells were naïve phenotype at both D15 and D26 timepoints suggestive of a lack of activation for even the small fraction of CD8^+^ T cells that infiltrate the CPT6 TME. In contrast, E0771 CD8^+^ T cells were predominantly of the central memory and effector memory phenotype, a balance that shifted slightly in favor of central memory cells and increased naïve cells at D26 compared to D15. In the CD4^+^ T cell compartment, effector memory subset dominated both CPT6 and E0771 TMEs at D15 timepoint suggesting that CD4^+^ T cells are the early responders to the developing tumor rather than CD8^+^ T cells. By D26, similar to CD8 T cell compartment, CD4 T cells were also predominantly central and effector memory phenotype in E0771 and naïve and central memory phenotype in CPT6 cells. Together, these data suggest a lack of activation signal for T cells in the CPT6 TME compared to E0771 TME as the tumor evolves.

We also analyzed the expression of inhibitory receptors PD-1, LAG3 and TIM-3 on the surface of CD8^+^ and CD4^+^ T cells infiltrating CPT6 and E0771 tumors (**Figure S4c and d**). In the CD8^+^ T cell compartment, we found low expression of PD-1, little to no expression of LAG3 and low to moderate expression of TIM3 in CPT6 at both timepoints (**Figure S4c**). In contrast, we found as high as 70% of E0771 tumor infiltrating CD8^+^ T cells to express PD-1 and >20% to express TIM3 at D15, which doubled to almost 40% by D26. In the CD4^+^ T cell compartment, a higher frequency of cells expressed PD-1 (∼50%) compared to CD8^+^ T cells in CPT6 (**Figure S4d**), providing further evidence of CD4 function in CPT6 TME. Similar to CD8^+^ T cell compartment, PD-1 and TIM3 expressing CD4 T cells dominated the two TMEs at both time points.

Given the importance of macrophages in breast tumor TME and their high proportion in CPT6 and E0771 tumors, we investigated their phenotype to gain insight into their potential role in the TME. Starting at D15, we found CPT6 tumors to harbor significantly more M2-like (CD206^+^) pro tumor and significantly lower M1-like (MHCII^+^) anti-tumor macrophages compared to E0771 tumors (**Figure 4h, upper panel**). By D26, both TMEs had close to double the percentage of CD206^+^ M2-like macrophages at D15 but maintained the significant difference between CPT6 and E0771 (**Figure 4h, lower panel).** Interestingly, while the proportion of M1-like macrophages decreased in CPT6 tumors from D15 to D26, that population was maintained in the E0771 tumors indicating a better pro and anti- tumor macrophage balance in the E0771 TME compared to CPT6 TME. These changes in the macrophage phenotypes were restricted to the tumor as macrophages in the spleen showed no significant difference between CPT6 and E0771 (**Figure 4h, right panels**). As anticipated, macrophages in the spleen were predominantly MHCII^+^ M1-like macrophages with little to no presence of M2-like macrophages (<5%) at D15 and D26. Additionally, we also investigated the presence of CX3CR1^+^ immunosuppressive macrophages in both TMEs and found CPT6 tumors to harbor significantly greater percentage of these at D15 and D26 (**Figure S4d**). Comparatively, less than 10% of CX3CR1 macrophages were found in the spleen at D15 which increased at D26 in CPT6 but not in E0771 demonstrating an enrichment of pro-tumor immunosuppressive macrophage phenotype in CPT6 tumors (**Figure S4e**).

## Discussion

In this study, we established CPT6, a novel syngeneic mouse ILC cell line derived from a spontaneous mammary tumor in a C57BL/6 mouse with dual knockout of *CDH1* and *PTEN*. To our knowledge, CPT6 is the first ILC-derived mouse cell line that is orthotopically transplantable into immune-competent syngeneic mice, addressing a critical gap in the field. In previously reported GEMM-derived models, the *TP53/CDH1* dual-knockout cell line requires an immunocompromised host for orthotopic tumor formation ^[10]^; the *CDH1/PIK3CA*-activation model did not yield transplantable cell lines ^[11]^; and the spontaneous *CDH1/PTEN* co-loss FVB/N model supported orthotopic growth only by tissue graft, not cell injection ^[12]^. CPT6 therefore enables the study of tumor growth, metastasis, and therapeutic response within an intact immune system, making it a uniquely valuable preclinical platform for immunotherapy and combination strategies.

CPT6 recapitulates key histological and molecular features of human ILC. The classical single-file growth pattern, complete loss of E-cadherin, and cytoplasmic redistribution of p120 confirm the defining phenotypic characteristics of this subtype. The spindle-cell and pleomorphic morphology, combined with a high mitotic rate, more closely resemble human pleomorphic ILC (pILC), a rare but aggressive subtype. Notably, in the previously described *CDH1/TP53* dual-knockout model on an FVB/N background, spontaneous tumors and derived cell lines were also described as pleomorphic ^[10]^, whereas the CDH1/PTEN FVB/N model displayed more classical ILC morphology ^[12]^. These differences suggest that genetic background and/or subclone selection during FACS sorting may contribute to morphological divergence across models. We observed that *CDH1* harbors a homozygous exon 6-10 deletion, which likely permits partial transcription yet abolishes functional protein. This highlights an important mechanistic nuance: exon-level genomic deletion does not necessarily eliminate transcription but can still abolish protein function — an event that may be underappreciated in analyses relying primarily on mRNA readouts. *PTEN* deletion sensitizes CPT6 cells to AKT pathway inhibition, supporting *PTEN* loss as a potential biomarker of capivasertib response. The intermediate sensitivity relative to PIK3CA H1047R-mutant BCK4 cells suggests that distinct PI3K/AKT alterations confer graded levels of AKT dependency and therapeutic vulnerability.

Transcriptomic profiling classifies CPT6 within the LumA subtype, consistent with the majority of human ILC and supported by positive CK8 staining in vitro. In vivo expression profiles also align more closely with LumA than with the luminal B model E0771, which exhibits higher proliferation and reduced reliance on canonical luminal programs. Beyond subtype classification, CPT6’s oncogenic mutation profile overlaps more closely with human ILC cell lines than with primary ILC tumors. The most clonally prominent driver mutations are Kras G12C and Cdh11 E462D. *CDH11* encodes type II cadherin, which facilitates cell-cell adhesion and epithelial mesenchymal transition ^[21]^; *CDH11* overexpression promotes breast cancer cell migration and bone metastasis ^[22,23]^. In CPT6, the E462D substitution likely has limited structural impact, yet the 100% variant allele frequency implies combined loss of heterozygosity and biallelic mutation, suggesting that loss of type II cadherin function may be a downstream consequence of *CDH1* knockout. The interplay between type I and type II cadherins warrants further investigation in ILC biology. The Kras G12C mutation — present in approximately 1.4% of ILC cases in TCGA -- expands the utility of CPT6 as a platform for testing Kras-targeted agents; CPT6 shows functional sensitivity to sotorasib, validating this mutation as a therapeutic target. In contrast, homozygous loss of *CDNK2A/B* did not significantly alter sensitivity to CDK4/6 inhibition, consistent with the concept that oncogenic driver activation (here, *KRAS*) can override tumor suppressor loss in determining therapeutic response ^[24]^. This raises the clinically relevant possibility that concurrent oncogene activation may limit CDK4/6 inhibitor monotherapy, warranting combinatorial strategies.

At the chromosomal level, CPT6 displays extensive copy number variation (CNV), including whole-chromosome gains and losses. This degree of chromosomal instability distinguishes CPT6 from most mouse mammary models, which are typically genomically stable despite clear oncogenic drivers ^[25]^, and increases its fidelity as a preclinical platform. The CNV pattern also resembles pILC, which more frequently harbors structural variants and CNV compared to classical ILC ^[6]^.

In vivo, CPT6 demonstrates rapid tumor growth consistent with the aggressive clinical behavior of pILC rather than classical ILC ^[7]^. Lung metastasis was consistently observed across multiple injection methods, with additional dissemination to ovaries and liver. While lung metastasis is less common in human ILC ^[26]^, reproducibility across experimental settings indicates a stable and tractable metastatic phenotype. Whether the *KRAS* G12C mutation, a canonical driver in lung adenocarcinoma, contributes to lung-homing is an important question for future study. Ovarian metastases were also observed, recapitulating a typical ILC metastatic site. Together, CPT6 offers a robust model for studying ILC metastasis, with rapid establishment of metastatic niches and retention of select ILC-relevant dissemination patterns.

The estrogen receptor biology of CPT6 warrants careful interpretation. ERα protein expression is low, yet the downstream ER response gene signature is enriched at levels comparable to LumA/B tumors, and functional assays confirmed ER pathway activity, including induction of *Greb1* and *Pgr*. These findings indicate that ER signaling is active in CPT6 despite low receptor protein levels. E2 treatment did not enhance proliferation (**Figures S3f, S3j**), a finding that should be interpreted cautiously given the rapid baseline growth kinetics and sensitivity to seeding density. The limited response to estrogen, fulvestrant, and their combination suggests that rapid intrinsic growth, driven by pleomorphic features and aberrant genomic alterations, overrides ER signaling. ERα overexpression also failed to restore endocrine sensitivity. Importantly, CPT6 captures luminal transcriptional identity but not classical endocrine sensitivity; this distinction must be considered when selecting this model for hormone-response studies.

A particularly valuable attribute of CPT6 is its ability to recapitulate the ILC immune microenvironment in an immune-competent host. CPT6 reveals a tumor microenvironment (TME) that is largely T cell-independent and dominated by myeloid populations. Tumor growth was comparable in wild-type and RAG-knockout mice, confirming that adaptive immunity plays a limited role in controlling tumor progression. CPT6 tumors are instead characterized by abundant granulocytic MDSCs, neutrophils, and macrophages skewed toward an M2-like, CX3CR1+ immunosuppressive phenotype. Temporal profiling at Day 15 and Day 26 reveals progressive deepening of immunosuppression as tumors grow. These features closely align with the immunosuppressive microenvironment observed in human ILC ^[9]^.

T cell infiltration in CPT6 tumors is low and functionally limited. The predominance of naive phenotypes and low expression of exhaustion markers suggest insufficient priming rather than classical checkpoint-mediated exhaustion, implying that immune evasion occurs primarily through T cell exclusion. CPT6 tumors also harbor a higher proportion of B cells (∼10%), consistent with the ER+ breast cancer immune landscape, although B cell enrichment has not been specifically reported in ILC versus NST and their prognostic significance remains undefined. The M2-macrophage and G-MDSC-dominated composition likely suppresses both T cell and B cell function despite the higher B cell proportion. Further investigation of B cell phenotypes and tertiary lymphoid structure formation in the CPT6 TME may clarify their role in ILC ^[9]^. Finally, plasmacytoid dendritic cells (pDCs) are significantly depleted in the CPT6 TME relative to E0771. Given the central role of pDCs in type I IFN secretion and adaptive immune activation, their absence is likely an additional contributor to the immunosuppressive state, even though pDC enrichment has not been specifically linked to ILC in human data ^[9]^. Together, these findings highlight the dominant role of myeloid cells in shaping the CPT6 TME, suggesting that strategies targeting the myeloid compartment, including macrophage reprogramming, CSF1R inhibition, or MDSC depletion, may be more effective than T cell-focused approaches alone.

This study has several limitations. CPT6 represents a single cell line and cannot capture the full heterogeneity of ILC. Its relatively rapid growth rate differs from the typically indolent clinical course of human ILC and may influence therapeutic response testing. The metastatic pattern, particularly predominant lung dissemination, does not fully recapitulate the clinical distribution of ILC metastases. Finally, low ERα expression and the absence of estrogen-dependent growth limit utility for endocrine therapy studies.

In conclusion, CPT6 is the first orthotopically transplantable, ILC-derived mouse cell line suitable for immune-competent syngeneic hosts. It recapitulates core molecular, histological, and immune features of human ILC, with resemblance to pILC; and provides a practical platform for evaluating immunotherapy, targeted therapy, and combination approaches. CPT6 is especially suited for interrogating the myeloid-rich, immunosuppressive TME that characterizes ILC and for testing innate immune-targeting agents. Future work will benefit from the development of additional syngeneic ILC models complementing CPT6, and from direct comparisons with GEMMs to delineate cell-intrinsic versus microenvironmental contributions to ILC biology in vivo.

---

**Figure S1.** Generation and cellular characterization of CPT6 cell line. **a.** Basic gating strategy for CPT6 cell sorting. FSC-A and SSC-A are used to gate alive cells, FSC-W and FSC-H are used to gate single cells. EpCAM+ cells are sorted as epithelial cells. EpCAM+ve cells in Epicult B culture are sorted via flow cytometry. **b.** Western blott confirming E-cad and PTEN knockout in CPT6 cell line. SSM3 WT cell line and SSM3 CDH KO with CRISPR-Cas9 are used as control. **c.** Cell cycle analysis of EpCAM+ve sorted cells. Cell cycle determined with FlowJo Dean-Jett-Fox analysis model.

**Figure S2.** In vitro and in vivo estrogen signaling response of CPT6. **a.** Western blot image showing expression of ERα detected by clone E115 (Millipore) antibody at low and high exposure in MCF-7, SSM3, CPT6 (passage 8 and 5), MM231 and 3T3 cell lines. **b.** Level of expression (kpm) of vital genes in CPT6 early passage and E0771 compared to ICLE data with denoted PAM50 subtypes (Basal, HER2, LuminalA, LuminalB). **c.** Relative fold change in expression of Esr1 (upper left) and Ppid (housekeeping gene) (upper right), mouse Greb1 (mGreb1, bottom left) and mouse PgR (mPgR, bottom right) in vehicle treated vs 1nM E2 treated E0771, CPT6 cell lines. **d.** Time course measurement of changing cell confluence as a result of hormone deprived CPT6 cell proliferation over a period of 72 hours in conditions like media containing (i) fetal bovine serum (FBS) (red) (ii) charcoal stripped serum (CSS) (blue) (iii) CSS+20pM E2 (green) (iv) CSS+ 1nM E2 (magenta) (v) CSS+1nM E2+ 100nM ICI (black). **e.** Estrogen response element (ERE) fluorescence assay of downstream ER signaling promoter activation in CPT6 and SSM3 with E2 and E2+ICI treatment. **f.** Bar graph depicting percent growth fold change by Presto Blue assay for CPT6 and SSM3 cells in vehicle (media) condition upon addition of E2 (1nM) alone, ICI (100nM) alone and E2 (1nM)+ICI (100nM). **g.** Experimental schematic to test impact of E2 on CPT6 tumor cells in vivo in B6 WT mice. **h.** Tumor growth curve showing tumor volume in mice injected with CPT6 cells (left panel) and E0771 cells (right panel) either without any circulating E2 (-E2) or supraphysiological levels of E2 (+E2). **i.** ESR1overexpression in CPT6 cell line (left); Maintenance of ESR1 overexpression in CPT6 cell line (right). **j.** E2 growth assay on CPT6 parental cell line, MCF7 cell line, EV and OE of CPT6 p5 in 10% CSS.

**Figure S3.** Molecular characterization of CPT6 transcriptome and genome. **a.** Scatter plot of somatic mutation variant allele frequency comparison in CPT6 cell line and B6 mouse liver. **b.** River plot of genomic subclone composition and driver oncogenes of each subclone in CPT6 revealed by Superfreq analysis. **c.** Reactome network of CPT6 somatic mutations. Arrow pointing to downstream pathways affected, top 50 reactome pathway nodes are shown. **d.** Tumor mutation burden comparison of CPT6 with human ILC cell lines (ICLE) and ILC samples in TCGA dataset (right). Tumor mutation burden (TMB) is calculated by mutations/Mb with maftool package.

**Figure S4.** Immune infiltration landscape of tumor microenvironment in CPT6 and E0771 in vivo tumor. **a.** Percentage of CD45+ cells in spleen and lymph node after CPT6 and E0771 cell injection. **b.** Flow cytometry workflow in identifying immune compositions in immune microenvironment analysis. **c.** Bar graphs depicting percentage of tumor infiltrating CD8 T cells expressing PD-1, LAG3 and TIM3 at D15 (left panel) and D26 (right panel) comparing CPT6 and E0771. **d.** Bar graphs depicting percentage of tumor infiltrating CD4 T cells expressing PD-1, LAG3 and TIM3 at D15 (left panel) and D26 (right panel) comparing CPT6 and E0771. **e.** Proportion of CX3CR1+ F4/80+ macrophages in tumor compared to spleen in CPT6 and E0771 at D15 (left panel) and D26 (right panel).

## Methods

### Mice and breeding strategy used in the study

To generate the Pten^L/L^. Cdh1^L/L^. Wap^Cre^ mice, Pten^L/L^ [B6.129S4-*Pten^tm1Hwu^*/J, Strain#006440, Jackson Laboratory], Cdh1^L/L^ [B6.129-Cdh1tm2Kem/J, Strain#005319, Jackson Laboratory] and WAP^cre^ [B6.Cg-Tg(Wap-Cre)11738MAM, Strain#01XA8 frozen embryos, NCI repository] were purchased/ obtained. Single flox Pten^L/L^, Cdh1^L/L^ were crossed with each other until homozygous double flox Pten^L/L^. Cdh1^L/L^ mice were obtained. These double flox mice were then crossed with WAP^Cre^ mice containing single copy of the transgene and bred until Pten^L/L^. Cdh1^L/L^ Wap^Cre^ ^+^ mice were obtained. Female mice were set up in breeding pairs with wildtype male or Pten^L/L^. Cdh1^L/L^ males to undergo at least 3 rounds of pregnancy for enhancing Cre recombinase activity and mammary epithelium specific Pten and Cdh1 deletion. Mice older than 8 months of age and/or 3 pregnancies were parked for observation of spontaneous tumor growth which was examined by palpating mammary glands on a weekly basis. C57BL/6J female mice purchased [B6/J, Strain#000664, Jackson Laboratories] were between 8-10 weeks of age. Animal protocols were approved by Institutional Animal Care and Use Committees (IACUC) of the University of Pittsburgh

### Generation of B6.ILC cell line and details of other cell lines used in the study

Spontaneous mammary tumor from an 8 months old Pten^L/L^.Cdh1^L/L^ Wap^Cre^ female mouse was harvested and digested under sterile conditions by mechanical disruption using scalpel followed by enzymatic digestion using a mix of collagenase IV (Worthington Biochemicals) and Dispase (Stemcell Technologies) both at 1mg/ml in Dulbecco’s modified Eagle’s medium (DMEM) media containing 10% fetal bovine serum (FBS) and 1% Pen/Strep. Resulting single cells were passed through 100um filter to remove tissue debris and cultured in Epicult B Mouse Mammary Epithelial Cell Medium Kit (Stemcell Technologies #05610). Cells were passaged 2-3 times to ensure outgrowth of adequate number of cells which were subjected to differential trypsinization for 2-3 passages to remove fibroblasts. Ultimately, epithelial cells were sorted by FACS process using EpCAM antibody. EpCAM positive live singlets were then cultured for 3-4 passages in Epicult B media supplemented with 10% FBS, 20ng/ml EGF (Stemcell Technologies), 100pM Estradiol (Sigma) and 1% Pen/Strep (Stemcell Technologies) (here on referred to as B6.ILC medium).

E0771 cell line was obtained from Dr. Robin Anderson (Olivia Newton-John Cancer Research Institute, Australia). Cells were cultured in DMEM (Gibco 11965-092) containing 20mM HEPES supplemented with 10% (FBS), and 1% Pen/Strep. SSM3 cell line was cultured in DMEM/F10 media supplemented with 10% FBS, 0.34% b-mercaptoethanol, 2.5mg/ml hydrocortisone, 2.5mg/ml transferrin and 10mg/ml insulin. B16-F10 melanoma cell line was obtained from Dr. M.J Turk (Dartmouth College, NH) and MC38 colon adenocarcinoma cell line was obtained from Dr. Jim P Allison (M.D. Anderson Cancer Center, TX). B16 cells were cultured in complete Roswell Park Memorial Institute (RPMI) media and MC38 in DMEM media both supplemented with 10% FBS, 1% Pen/Strep, 2mM glutamine, 1mM pyruvate, 5mM HEPES and 100μM non-essential amino acids. Human breast cancer cell line MCF7 was purchased from ATCC (cat # HTB-22) and cultured in DMEM with 10% FBS. All cell lines were cultured and maintained at 37C, 5% CO_2_. Cell lines were also periodically tested for mycoplasma and ensured to be clean of any contamination

### Immunohistochemical (IHC) staining

Mouse mammary tumors harvested from euthanized mice were divided into pieced prior to tissue dissociation and some pieces were formalin-fixed in 10% neutral buffered formalin (NBF) for 48 hours, embedded in paraffin, sectioned and stained with hematoxylin and eosin (H&E)per standard protocol. For assessing the expression of markers like E-cadherin, p120, Pten and Ki67, cell pellets from CPT6 and SSM3 cell lines were obtained after washing the cells with PBS by centrifugation at 500g for 5mins at 4C immediately followed by fixing in 10% NBF for 24hrs and transferred to 70% ethanol prior to embedding into formalin. FFPE cell pellet sections were cut at a thickness of 4 μm and were mounted on plus slides. Staining was performed using standard IHC protocol involving dehydration at 65 °C overnight, deparaffinization using xylene and rehydration with ethanol. Heat-induced epitope retrieval was performed using citrate buffers followed by 1hr of blocking before incubation with primary antibodies for 1 hour at RT or overnight at 4C. Secondary antibody–horseradish peroxidase was added after washing off unbound primary antibodies using Tris-buffered saline containing 0.1% Tween 20. Slides were then incubated with DAB chromogenic substrate (Abcam, cat# ab64238) and monitored visually for color development followed by washing with cold water. Finally, cells were counterstained with DAPI/ Hematoxylin for nuclear staining, mounted with mountant and stored at 4 °C in the dark until imaging. All antibodies, their catalog numbers and dilutions are listed in Supplementary Material 01.

### Quantification of IHC positivity

QuPath v0.6.0^[27]^ on Pitt HTC cluster was used to quantify positivity of Ki67% in IHC imaging. Hematoxylin OD mode was applied with nucleus parameters as followed: 15px background radius, 3px sigma, 10px^2 minimum area and 1,000px^2 maximum area. Intensity threshold were set as 0.1 and max background intensity as 2 and split by shape. Cell expansion was defined as 5px with cell nucleus included. DAB OD mean was set as compartment score. For intensity threshold, 1+ parameter was set as 0.2, 2+ parameter as 0.4 and 3+ parameter as 0.6 with single threshold mode.

### Immunofluorescence (IF) staining and imaging

3x10^5^ CPT6, MCF-7 cells were plated onto glass coverslips overnight and fixed using cold methanol after removing media and washing with PBS. Cells fixed onto the coverslips were then washed to remove residual methanol prior to transferring to a humidity chamber for staining. Cell were blocked using blocking buffer (PBS+5%BSA+0.3% Triton X100) and incubated with primary antibodies for Cytokeratin 8 (Abcam cat # ab53280) at 1:1000 and E-cadherin (BD, cat #610182) both at 1:100 dilution overnight at 4C. Primary antibodies were then washed off and secondary antibodies {goat-anti-mouse-AF488 and goat-anti-rabbit-AF647, (Thermo Scientific Cat # A11001 and A21245)}, both at 1:200 were incubated for 1hr at room temperature prior to washing and staining with Hoechst dye (1:200 in PBS) for nuclear staining. Coverslips were then washed and mounted onto slides for imaging which was performed on a fluorescence microscope (Olympus, CZX16) at 20x magnification to capture images. All antibodies, their catalog numbers and dilutions are listed in Supplementary Material 01.

### Western blot for estrogen receptor alpha expression

Total protein was extracted from CPT6, MCF-7, MM231, SSM3 and 3T3 cell lines by lysing at least 5x10^5^ cells already in culture at least 12hrs by using 5% sodium dodecyl sulfate (SDS) buffer and sonicated. Total protein was then quantified by BCA assay and 40ug was loaded onto SDS polyacrylamide gel and resolved through electrophoresis. The proteins were then transferred onto nitrocellulose membrane for immunoblotting. First, the membrane was blocked for non-specific binding using blocking buffer (Intercept Protein-free blocking buffer, Li-Cor cat#927-90003). Primary antibodies to probe expression of ERa (Millipore cat # 04-820, Clone 60C) and beta actin as a loading control (Sigma Aldrich, cat#A5441) were both used at 1:2000 dilution. Secondary antibodies, goat anti-Rabbit IgG (H + L), (Licor cat# 926-32211) for ERa and goat anti-mouse IgG (H+L) (Licor cat# 925-68020) for actin were added to the blot and imaged with two different exposure settings to capture lighter ERa bands. All antibodies, their catalog numbers and dilutions are listed in Supplementary Material 01.

### Cell cycle analysis using Propidium iodide (PE)

CPT6 cells were fixed using cold 70% ethanol and incubated at 4C for 30mins prior to washing 3 times by centrifugation at 500g for 5mins with PBS to remove excess ethanol. Cells were stained with 0.5ml of FxCycle PI/RNase staining solution (Thermo Fisher, cat # F10797), incubated for 30mins at room temperature in the dark and immediately acquired on flow cytometer. Data was analyzed using manual gating and automated cell cycle analysis module in FlowJo software (BD)

### Bulk RNA sequencing of cell lines and tumor

Bulk RNA sequencing was performed on CPT6 cells at two different cell passages (to account for potential changes *in vitro* during culturing) and from FACS sorted live EpCAM+ tumor cells isolated from *in vivo* tumors harvested at day 12 post injection with 5x10^5^ CPT6 and E0771 injection into the mammary fat pad of WT B6 female mice. Briefly, tumors were resected, digested into single cells using collagenase and dispase, stained with viability dye and EpCAM antibody and sorted using flow cytometry. For both *in vitro* cell lines and cells isolated from D12 tumors, RNA was extracted by lysing cells in buffer containing RNase inhibitor (Clontech) and 0.2% Triton X-100 (Sigma Aldrich). Once completed, cells underwent reverse transcription (Applied Biosystems RT Kit) followed by a cDNA amplification and then bead selection to target fragment sizes around 1.5-2kb.cDNA was then diluted to 0.2ng/ul and carried into the library construction step using i7 and i5 Index primers (Invitrogen i7 Barcoding Primers and Invitrogen i5 Barcoding Primers). After the final library was constructed, the fragment size was quality checked by TapeStation 500 and quantified using a Qubit. All libraries were sequenced on an Illumina NextSeq™ 500/550 High Output v2 kit for 75 cycles at the UPMC Children’s Sequencing Core with a sequencing depth of at least 40 million reads/library. Raw single-end FASTQ files were generated using Illumina sequencing platforms with base-calling and demultiplexing performed by bcl2fastq version 2.20. Read quality was assessed using FastQC version 0.11.9, followed by adapter trimming and quality filtering with Trimmomatic version 0.39 using TruSeq3 adapters. Trimmed reads were aligned to the mouse reference genome (mm10) using HISAT2 version 2.1.0, and alignments were sorted with samtools version 1.15.1. Gene-level read counts were quantified using HTSeq-count version 0.13.5. All three datasets (SET1–SET3) were processed using this identical workflow. Detailed information of 3 sets of RNAseq is described in Supplementary Material 02.

The analysis includes PAM50 breast cancer subtype classification using the genefu package, with samples compared to four external datasets: (1) GSE223630 public mouse tumor RNA-seq data^[28]^ for cross-study validation, (2) ICLE breast cancer cell line panel for passage stability assessment ^[29]^, (3) TCGA (The Cancer Genome Atlas) breast cancer samples and (4) PDO sequencing dataset of breast cancer in described by Sachs *et.al* ^[30]^ for clinical subtype comparison. Key analytical approaches include batch effect correction using ComBat (sva package), quantile normalization (preprocessCore), dimensionality reduction via t-SNE (Rtsne) for visualization of sample relationships, differential gene expression analysis (edgeR), and pathway enrichment analysis using GSVA with MSigDB Hallmark gene sets (GSVA, msigdbr packages). Mouse genes are converted to human orthologs using the Mouse Genome Informatics homology database to enable cross-species comparisons. Original data can be downloaded from GEO through submission number GSE316819.

### Whole exome sequencing

CPT6 cells passage 33 and paired B6 mouse liver parenchymal cells had DNA extracted with Zymo Research quick-DNA microprep Kit and sent for sequencing with Twist Mouse Exome Panel from Twist Bioscience. Library preparation was performed according to the manufacturer’s guidelines. The sequencing libraries were clustered on a 25B flow cell on the Illumina NovaSeq X Plus instrument. Preprocessing of sequencing files including quality control, adapter trimming and genomic alignment were performed following GATK best practice. For oncogene visualization, we used the vep/95 annotation .txt output and retained variants with moderate or high functional impact. Mouse gene names and human homologs were added through the Ensembl API. Using the OncoKB oncogene list from MSKCC ^[31]^, we prioritized high-impact mutation classes to generate oncoplots and mutation-type bar plots for CPT6. We further compared SBS mutational signatures between CPT6 and human ILC cell lines from the ICLE project using SigProfilerAssignment. For tumor mutation burden (TMB), VCFs were converted to MAF using vcf2maf/1.6.8, then filtered following the ICLE pipeline: removal of non-protein-coding genes, variants outside coding regions, variants with AF < 0.1 or DP < 10, genes without human homologs, and silent/intronic mutations. TMB was then computed from exome-restricted MAF files using maftools. For CNV analysis, we used CNVkit/0.9.5 generated .cnr files, removed antitarget regions, retained OncoKB genes, and calculated gene-length-weighted log2(tumor/liver) values. We verified that weighting was appropriate by showing no correlation between gene length and log2 ratio. CNVs were called using ±0.3 thresholds, with log2 < -1.17 defining homozygous deletion. Heatmaps were generated for non-homozygous CNVs, while homozygously deleted genes (*CDH1, PTEN, CDKN2A, CDKN2B*) were plotted separately. Samtools/1.22.1 bedcov was used to determine exon loss status. Gain/loss gene sets were analyzed by gseapy in Python 3. For subclone and driver analysis, SuperFreq was run from tumor/liver Mutect2 VCFs, BAMs, BED, and reference FASTA. SuperFreq-derived somatic mutations were reanalyzed through the oncogene pipeline, and CNV patterns were concordant with CNVkit/Control-FREEC/11.5. Finally, we performed Reactome-based pathway analysis on non-synonymous somatic genes with human homologs and built an interactive vis.js pathway network with data from NCBI2Reactome pathway. (<u>Home - Reactome Pathway Database</u>). Mutation frequency was compared with TCGA dataset in Thennavan et. al 2021^[19]^. Original data of WES sequencing can be downloaded from SRA through submission number 15983533. Somatic and oncogenic mutation are summarized in Supplementary Material 03.

### Dose response assay

CPT6 cells were seeded at 3,000 cells/well in 24-well plates with 500 μL complete medium (DMEM + 10% FBS + 1% antibiotic-antimycotic) and cultured overnight. Other cell lines were seeded at 5,000 cells/well in 96-well plate with 100uL medium. Abemaciclib (CDK4/6 inhibitor), sotorasib (KRAS G12C inhibitor) and capivasertib (AKT inhibitor) were dissolved in DMSO to 10 mM stock solutions. On Day 2, compounds were serially diluted to a starting concentration of 6×10⁴ nM. 100 μL of diluted compound (6× concentration) was added to wells containing 500 μL medium to achieve a final starting concentration of 10,000 nM. Control wells received 100 μL medium only. Treatments were performed in duplicate. On Day 6, cell viability was assessed using PrestoBlue (Thermo Fisher) reagent. 66 μL PrestoBlue (10% v/v) was added to each well and incubated for 2 hours. Fluorescence was measured at 570 nm excitation. Cell-free controls (medium + PrestoBlue, no cells) were prepared in parallel. Viability was calculated as: Viability (%) = [(Sample fluorescence - Cell-free control) / (Untreated control - Cell-free control)] × 100. For other cell lines, drugs were added to same concentration gradient to total of 200 μL medium. 22 μL of Presto Blue reagent was added to measure the fluorescence. Dose-response curves were analyzed using three-parameter logistic regression in GraphPad Prism 10. F-test was performed with null hypothesis that one curve will fit for all groups of data.

### Quantitative reverse transcription PCR (qRT-PCR)

Cell lines were seeded in triplicates into 6-well plates with 100,000 cells per well, respectively. RNA was extracted after desired treatments and then synthesized to cDNA using PrimeScript RT Master Mix kit (#RR036, Takara Bio). qRT-PCR reactions were performed with SybrGreen Supermix (catalog no. 1726275, Bio-Rad), and the ΔΔ*C*_t_ method was used to analyze relative mRNA FCs with *RPLP0* as the internal control. All primer sequences are indicated below:

*Esr1*(forward): 5’-TCTGCCAAGGAGACTCGCTACT-3’

*Esr1*(reverse): 5’- GGTGCATTGGTTTGTAGCTGGAC-3’

*Ppid*(forward): 5’- GCTTAGAGGCTCTTGAAATGGA-3’

*Ppid*(reverse): 5’- GAAACACAACAGGGAGCATGTA-3’

*Greb1*(forward): 5’- CGCTCTTTGTCCTCATCCACGA-3’

*Greb1*(reverse): 5’- GTTGAGCAGTCTGTCCACTAACC-3’

*PgR*(forward): 5’- TCGCCTTAGAAAGTGCTGTC-3’

*PgR*(reverse): 5’-GCTTGGCTTTCATTTGGAACG-3’

Statistical analyses were performed with t-test and Holm-Sidak correction, with p<0.05 as statistical significance cutoff.

### Estrogen Receptor Element (ERE) reporter assay

CPT6 and SSM3 cells were seeded in replicates of 6 in 96-well-plates at 7,480 cells/well and 18,700 cells/well, respectively. Upon reaching 70% confluency at approximately 24 hours, cells were incubated at 37C, 5% CO2 in 100µl PBS supplemented with Mg2^+^ and Ca2^+^ for 45 minutes. Media was aspirated and replaced with 100µl modified improved MEM with 10% charcoal stripped serum (CSS), 1% Pen/Strep, 20ng/ml EGF (CPT6) or 100µl 10% CSS phenol free DMEM/F12, 500µl 0.34% b-mercaptoethanol, 2.5mg/ml hydrocortisone, 2.5mg/ml transferrin, 10mg/ml insulin (SSM3). Following 24 hours of hormone deprivation, cells were co-transfected at a ratio of 5:1 with a vector harboring a firefly luciferase gene under an estrogen-response-element promoter (ERE-Tk-Luc, 100ng) and a vector harboring Renilla luciferase under a null promoter (20ng) using the Lipofectamine 3000 (Invitrogen) method. 24 hours post-transfection, cells were either treated with vehicle control (EtOH), 1nM 17b-estradiol (E2, Sigma-Aldrich, Cat#E2257) or E2 +100nM Fulvestrant (ICI182,780, Tocris Bioscience Cat #1047) for 24 hours. Cells were lysed and assayed using the Dual-Glo kit following the manufacturer’s protocol (Promega #E2940). Luminescence was measured with GLOMAX®-Multi+ Microplate Multimode Reader (Promega). All firefly luciferase values were normalized to non-transfected control cells and the Renilla luciferase transfection control. The fold change in luciferase activity for each cell line with comparted to the respective vehicle treated cells and is presented in bar graphs with the standard error of the mean. Statistical analysis was performed using a one-way ANOVA with Dunnett’s post hoc test for multiple comparisons

### Cell proliferation assay and hormone deprivation

To test the impact of estradiol hormone on CPT6 cells, we first deprived the cells of the hormone by culturing them in IMEM media containing 5% Charcoal stripped serum (CSS) and incubated at 37C, 5% CO_2_. This CSS containing media was refreshed twice a day to ensure removal of debris and effective hormone deprivation. At D3, cells were trypsinized and added to 96 well plates at 2000 cells/ well. Cells were treated as indicated with (1) media containing FBS, (2) media containing CSS (3) media containing CSS+ 20pm E2 (4) media containing CSS+ 1nM E2 (3) media containing CSS+ 1nM E2+ 100nM ICI (4) media containing CSS+ 10nM E2 (5) media containing CSS+ 100nM ICI. Proliferation was tested by (a) continuous monitoring of cell growth and confluence for 72hrs in the wells measured as phase contrast with the Incucyte Live cell imaging system. (b) flow cytometric analysis through cell trace violet (CTV) dye labeling (Invitrogen Cat # C34571) prior to incubation for 24 hr, 48 hr and 96 hrs (c) PrestoBlue fluorescent reagent (Invitrogen Cat # A13261) assay following manufacturers protocol to measure cell viability after adding presto blue to wells and incubating at 37C for 10mins

### Testing the impact of estradiol on CPT6 tumor growth *in vivo*

To test *in vivo* the impact of estradiol on the development and growth of implanted CPT6 cells, we utilized a cohort of WT B6 mice between the age of 6-10 weeks and used E0771 cell line as a control. First, to remove the source of circulating estrogen, ovariectomy surgery was performed (D-16) by anaesthetizing mice via isoflurane. Lower dorsal area was shaved, and an incision was made to remove ovaries after cauterizing the blood vessels. Wound clips were used to close the surgical incision, and mice were treated with opioid analgesic buprenorphine for pain management for 2-3 days. Post-surgery on D-2, mice were divided into two groups- one with E2 pellets implanted subcutaneously at the back of the neck using a 12-gauge trocar to achieve supraphysiological levels of circulating E2 and other set underwent sham implantation. Both groups of mice were then injected on D0 with 5x10^5^ cells (CPT6 or E0771) into the mammary fat pad and tumor growth was measured for 30 days using digital calipers. Tumor volume was calculated as [(width)^2^ x length/ 2] was calculated blinded and randomized

### Tumor cell injections and growth kinetics

Female mice between 8-10weeks of age were used for all experiments. For CPT6 and E0771 tumors, 5x10^5^ cells were injected subcutaneously into the mammary fat pad using a 30-needle gauge insulin syringe. Mice were anaesthetized briefly using isoflurane prior to injection of cells. Tumors were measured every 3 days starting D9 using digital calipers and tumor volume [(mm^3^), calculated as (width)^2^ x length/ 2] was calculated blinded and randomized. Mice were euthanized at indicated time points for study or when the tumor reached a mean diameter of 15mm.

### Flow cytometry for immune infiltration analysis

At experimental end points, mice were euthanized using approved methods of CO_2_ chamber and secondary cervical dislocation and immediately dissected to harvest implanted CPT6 and E0771 tumors along with spleens and non-draining axial/ thoracic lymph nodes. Samples were collected in cold DMEM+ 2% FBS processing media on ice to prevent tissue degradation and cell death. Tumor processing was performed by mechanical and enzymatic methods as described above. Spleens were mechanically crushed using glass slides in a petri dish and cells were filtered through 40um filters to remove tissue debris. Tumor and spleen single cell suspensions were washed using processing media and RBC lysis was performed to remove red blood cells using 1x RBC lysis buffer (Invitrogen Cat # 00-4333-57). Samples were then washed with FACS buffer (PBS+ 2.5% BSA) incubated with surface antibodies (listed in **Supplementary Material 01**) for 30mins on ice in the dark. Cells were washed by centrifugation at 500g for 5mins and fixed using eBioscience Foxp3/ Transcription Factor Staining Buffer Set (Invitrogen, cat#00-5523-00) for 15 mins at 4C and washed again with 1x permeabilization buffer from the kit. Intracellular staining for Foxp3 was performed for 30min at room temperature in 1x perm buffer, and cells were washed and resuspended in FACS buffer until acquisition on Aurora Cytek spectral flow cytometer. Downstream analysis was performed using FlowJo (TreeStar/BD, v10.1). All antibodies, their catalog numbers and dilutions are listed in Supplementary Material 01.

### Mammary Fat Pad Clearing and Injection

Female mice (3-4 weeks old) were anesthetized with 2-2.5% isoflurane in oxygen and placed supine on a surgical platform with continuous isoflurane delivery. The surgical area was disinfected with 70% ethanol followed by betadine. Two subcutaneous incisions were made: one along the midline (∼1.5 cm) and one on each hindleg (∼1.2 cm). The inguinal (#4) mammary fat pad was exposed using sterile cotton swabs. The lymph node was identified as a central landmark, and three converging blood vessels were cauterized. The mammary epithelium was cleared by cutting just outside the lymph node (nipple side) and across the fat pad width adjacent to the lymph node. Adjacent #3 and #5 mammary tissue was removed to prevent invasion. The cleared #4 fat pad was separated laterally and from underlying skin, remaining attached only at the body cavity side. Cell suspensions (in 10 μL media) were prepared and kept on ice. Using a 25 μL Hamilton syringe with 26-gauge needle, 10 μL of cell suspension was injected into the cleared fat pad. Successful injection was confirmed by visible bubble formation. Incisions were closed with 4 wound clips (removed at 7-10 days). Mice recovered on a heating pad (38°C) and were monitored daily for 3-5 days post-surgery. Tumor growth was measured weekly with calipers; mice were euthanized if tumors reached 2 cm diameter or ulcerated.

### In Vivo Bioluminescence Imaging (IVIS)

Protocol is adapted from Chen *et. al* ^[32]^. An IVIS Lumina X5 system (Perkin Elmer) was used to evaluate metastatic disease burden by bioluminescent imaging of mice weekly. The disseminated CPT6 RFP/Luc cells were detected by injection of a luciferase substrate: (D-Luciferin (Cat No: LUCK-1G; Gold Biotechnology), resuspended in sterile PBS (Ca*2+* or Mg*2+* free) and injected IP at a dose of 150 mg/kg body weight. Ten to fifteen minutes after luciferin injection, mice were anesthetized with isoflurane/oxygen and positioned on a warmed stage in the IVIS imaging chamber to expose the mammary fat pad and ventral body surface. Imaging was conducted using consistent image settings for all mice (height, binning, FStop) and exposure time (0.25–60s). Bioluminescence signal was reported using Living Image software (Caliper Life Sciences).

### Mouse mammary intraductal (MIND) injection

MIND injections were performed according to Sflomos *et al.* ^[33]^ and Behbod FS *et al.* ^[34]^ without surgically opening the mouse. In brief, mice were anesthetized with isoflurane, and the inguinal nipples were prepared for injection under a stereomicroscope. Hair surrounding the target nipple area was removed using Nair depilatory cream, and any dead skin or debris covering the nipple opening was cleared using fine micro-dissecting tweezers. A 5-10 µL aliquot of the CPT6 cell suspension was loaded into a 25 µL syringe fitted with a 33-gauge metal-hub needle. The nipple was held gently with the fine tweezers and lifted slightly to align with the injection needle. The needle tip was inserted into the nipple opening, and the solution was slowly injected at a controlled rate of approximately 40 ul/minute to minimize potential damage to the ductal lumens from rapid flow. Injection was performed once per nipple.

## Supplementary materials

**Supplementary 01: Antibody information**

**Supplementary 02: RNAseq metadata**

**Supplementary 03: WES metadata**

**Supplementary 04: Figure 1 raw data**

**Supplementary 05: Figure 2 raw data**

**Supplementary 06: Figure 3 raw data**

**Supplementary 07: Figure 4 raw data**

**Supplementary 08: Figure S1 raw data**

**Supplementary 09: Figure S2 raw data**

**Supplementary 10: Figure S3 raw data**

**Supplementary 11: Figure S4 raw data**

## Data availability

Raw sequencing data of RNAseq is uploaded on GEO through submission number GSE316819. Raw sequencing data of WES is uploaded on SRA through submission number 15983533. Computational scripts for data processing and plotting are submitted on github: https://github.com/leeoesterreich/CPT6_code_manuscript. Cleaned RNAseq data of public breast cancer dataset is named ilc_clean_data.Rdata and uploaded together in Supplementary Material 05.

## Authors’ Contributions

### Conception and design

S. Onkar, A.V. Lee, D.A.A. Vignali, S. Oesterreich

### Development of methodology

S. Onkar, D. Liu, D. Seachrist, K.W. Bonk, L. Klei, J. Chen, K. Ding, L. Savariau. M. Yates, A.V. Lee, D.A.A. Vignali, S. Oesterreich

### Acquisition of data

(performed experiments, provided animals, cell lines, plasmids and other technical support): S. Onkar, D. Liu, D. Seachrist, C. Merkel, L. Klei, J. Chen, K.W. Bonk, K. Ding, L. Savariau. M. Yates, J. Hooda, L. Stabile, K. Ruth, C. Workman, A.V. Lee, D.A.A. Vignali, S. Oesterreich

### Analysis and interpretation of data

(statistical analysis, biostatistics, computational analysis): S. Onkar, D. Liu, D. Seachrist, J. Zou, L. Klei, J. Chen, K.W. Bonk, I. Thale, A.C.Chang, K. Ding, L. Savariau, M. Yates, A.V. Lee, G. Tseng, L Rigatti, P Lucas, D.A.A. Vignali, S. Oesterreich

### Writing, review, approval of final manuscript

S. Onkar, D. Liu, D. Seachrist, J. Zou, C. Merkel, I. Thale, A.C.C Chang, L. Klei, J. Chen, K.W. Bonk, K. Ding, L. Savariau, M. Yates, J. Hooda, L. Stabile, L. Rigatti, P.C. Lucas, G. Tseng, K. Ruth, C.J. Workman, A.V. Lee, D.A.A. Vignali, S. Oesterreich

### Study supervision

A.V. Lee, D.A.A. Vignali, S. Oesterreich

## Acknowledgement

We thank Dr. Osama Shiraz and Dr. Fangyuan Chen for providing support to access and analysis of ICLE data. Daisong Liu was a visiting research scholar at the University of Pittsburgh School of Medicine supported by funds from The China Scholarship Council and Tsinghua University.

## Disclaimer

During the preparation of this work the author(s) used Claude (Anthropic) to improve the clarity and phrasing of the text in English. After using this tool/service, the author(s) reviewed and edited the content as needed and take(s) full responsibility for the content of the publication.

