## Supplementary material for "Novel transplantable mouse cell line model recapitulates invasive lobular breast carcinoma (ILC) phenotype and immune microenvironment": Suppl figures

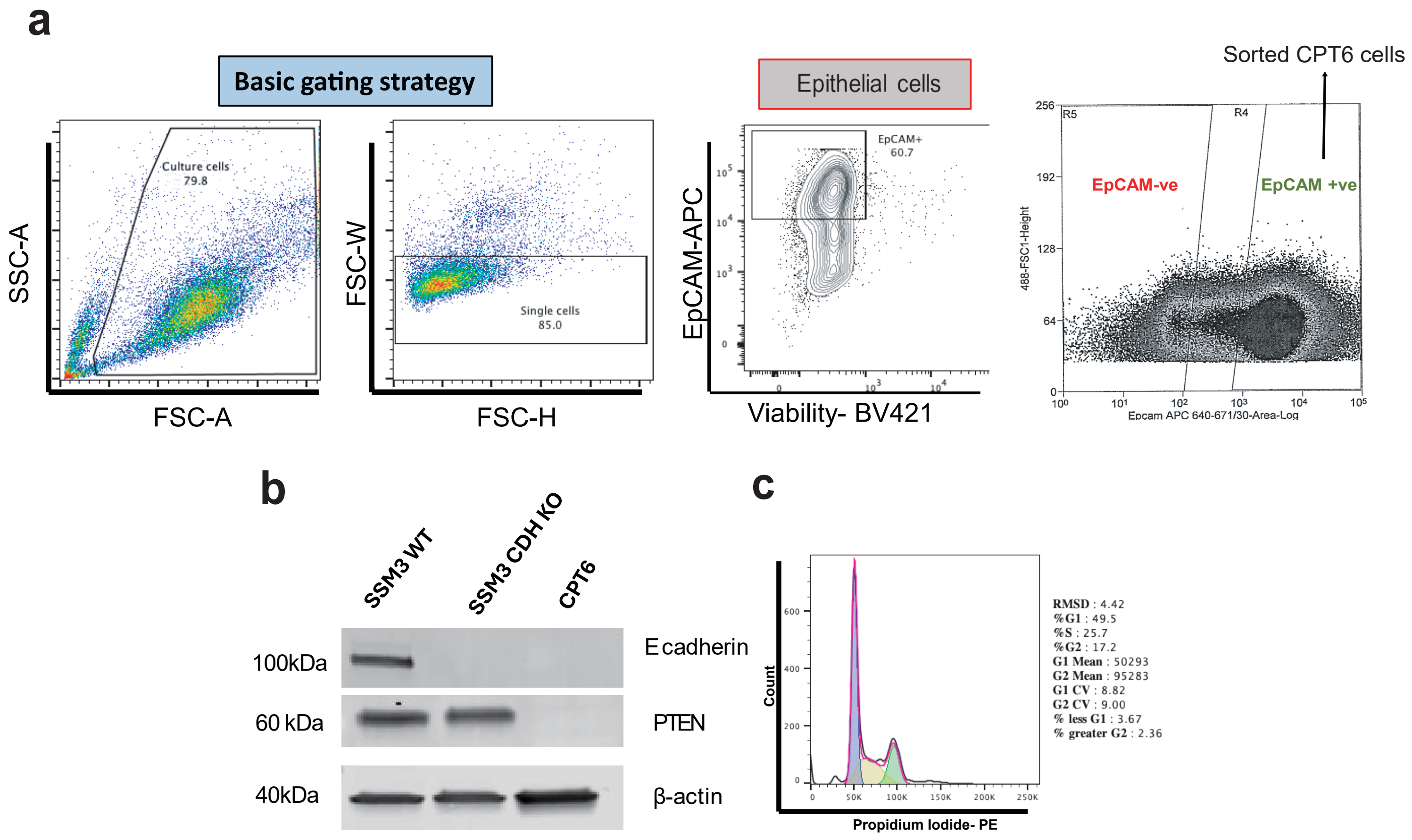


**Figure S1. Generation and cellular characterization of CPT6 cell line. a.** Basic gating strategy for CPT6 cell sorting. FSC-A and SSC-A are used to gate alive cells, FSC-W and FSC-H are used to gate single cells. EpCAM+ cells are sorted as epithelial cells. EpCAM+ve cells in Epicult B culture are sorted via flow cytometry. **b.** Western blott confirming E-cad and PTEN knockout in CPT6 cell line. SSM3 WT cell line and SSM3 CDH KO with CRISPR-Cas9 are used as control. **c.** Cell cycle analysis of EpCAM+ve sorted cells. Cell cycle determined with FlowJo Dean-Jett-Fox analysis model.

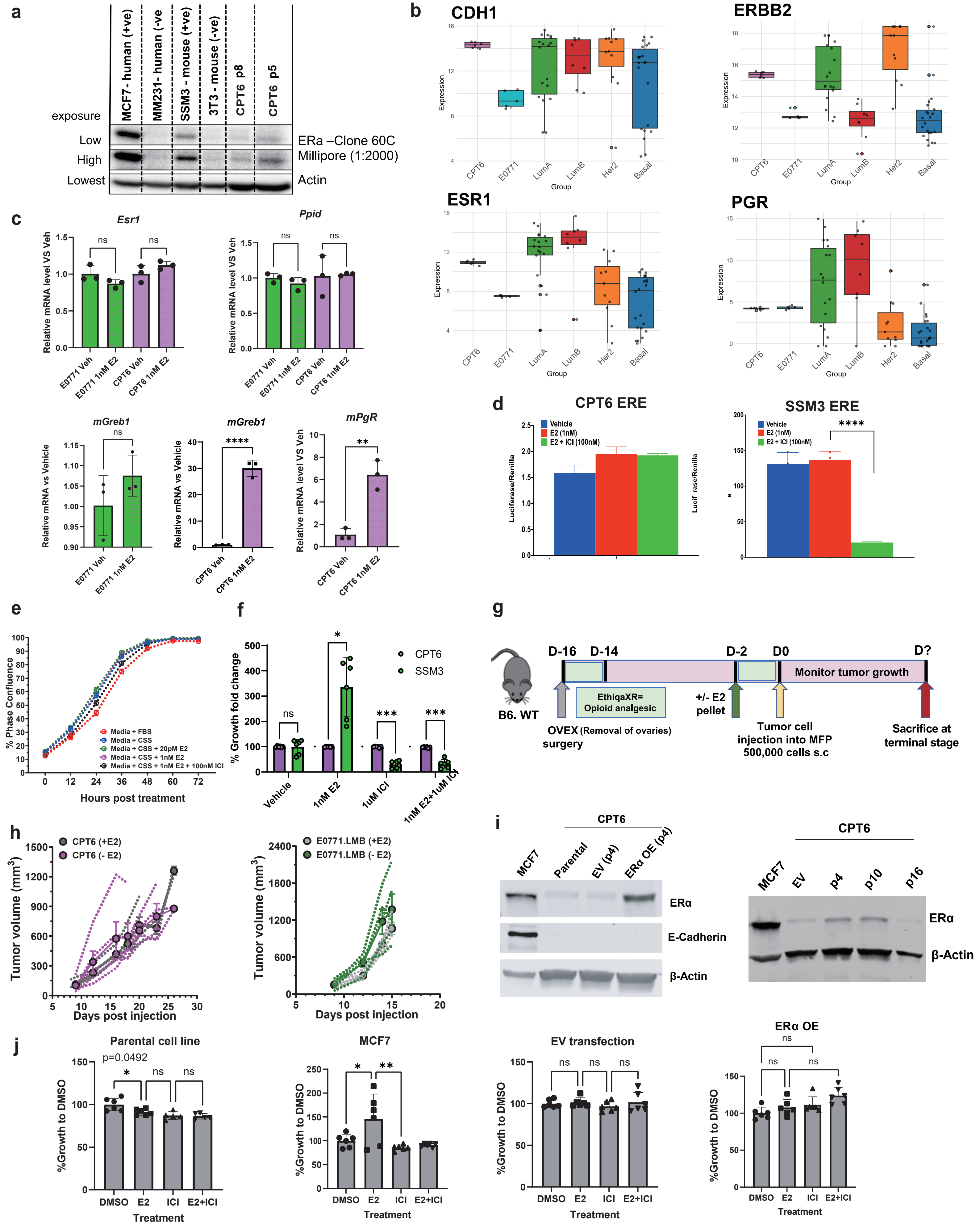


**Figure S2. *In vitro* and *in vivo* estrogen signaling response of CPT6. a.** Western blot image showing expression of ERα detected by clone E115 (Millipore) antibody at low and high exposure in MCF-7, SSM3, CPT6 (passage 8 and 5), MM231 and 3T3 cell lines. **b.** Level of expression (kpm) of vital genes in CPT6 early passage and E0771 compared to ICLE data with denoted PAM50 subtypes (Basal, HER2, LuminalA, LuminalB). **c.** Relative fold change in expression of Esr1 (upper left) and Ppid (housekeeping gene) (upper right), mouse Greb1 (mGreb1, bottom left) and mouse PgR (mPgR, bottom right) in vehicle treated vs 1nM E2 treated E0771, CPT6 cell lines. **d.** Time course measurement of changing cell confluence as a result of hormone deprived CPT6 cell proliferation over a period of 72 hours in conditions like media containing (i) fetal bovine serum (FBS) (red) (ii) charcoal stripped serum (CSS) (blue) (iii) CSS+20pM E2 (green) (iv) CSS+ 1nM E2 (magenta) (v) CSS+1nM E2+ 100nM ICI (black). **e.** Estrogen response element (ERE) fluorescence assay of downstream ER signaling promoter activation in CPT6 and SSM3 with E2 and E2+ICI treatment. **f.** Bar graph depicting percent growth fold change by Presto Blue assay for CPT6 and SSM3 cells in vehicle (media) condition upon addition of E2 (1nM) alone, ICI (100nM) alone and E2 (1nM)+ICI (100nM). **g.** Experimental schematic to test impact of E2 on CPT6 tumor cells in vivo in B6 WT mice. **h.**Tumor growth curve showing tumor volume in mice injected with CPT6 cells (left panel) and E0771 cells (right panel) either without any circulating E2 (-E2) or supraphysiological levels of E2 (+E2). **i.** ESR1overexpression in CPT6 cell line (left); Maintenence of ESR1 overexpression in CPT6 cell line (right). **j.** E2 growth assay on CPT6 parental cell line, MCF7 cell line, EV and OE of CPT6 p5 in 10% CSS.

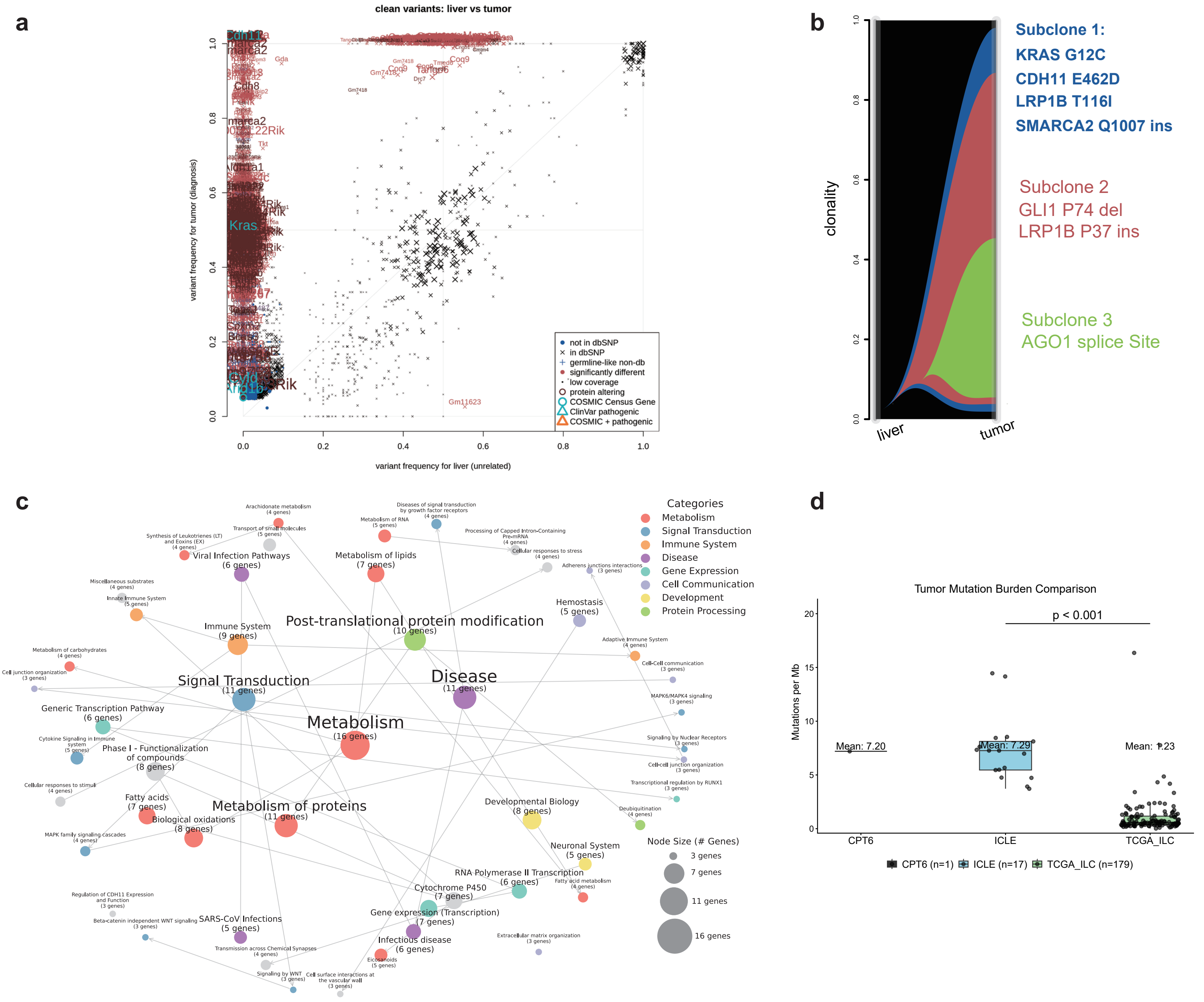


**Figure S3. Molecular characterization of CPT6 transcriptome and genome. a.** Scatter plot of somatic mutation variant allele frequency comparison in CPT6 cell line and B6 mouse liver. **b.** River plot of genomic subclone composition and driver oncogenes of each subclone in CPT6 revealed by Superfreq analysis. **c.** Reactome network of CPT6 somatic mutations. Arrow pointing to downstream pathways affected, top 50 reactome pathway nodes are shown. **d.** Tumor mutation burden comparison of CPT6 with human ILC cell lines (ICLE) and ILC samples in TCGA dataset (right). Tumor mutation burden (TMB) is calculated by mutations/Mb with maftool package.

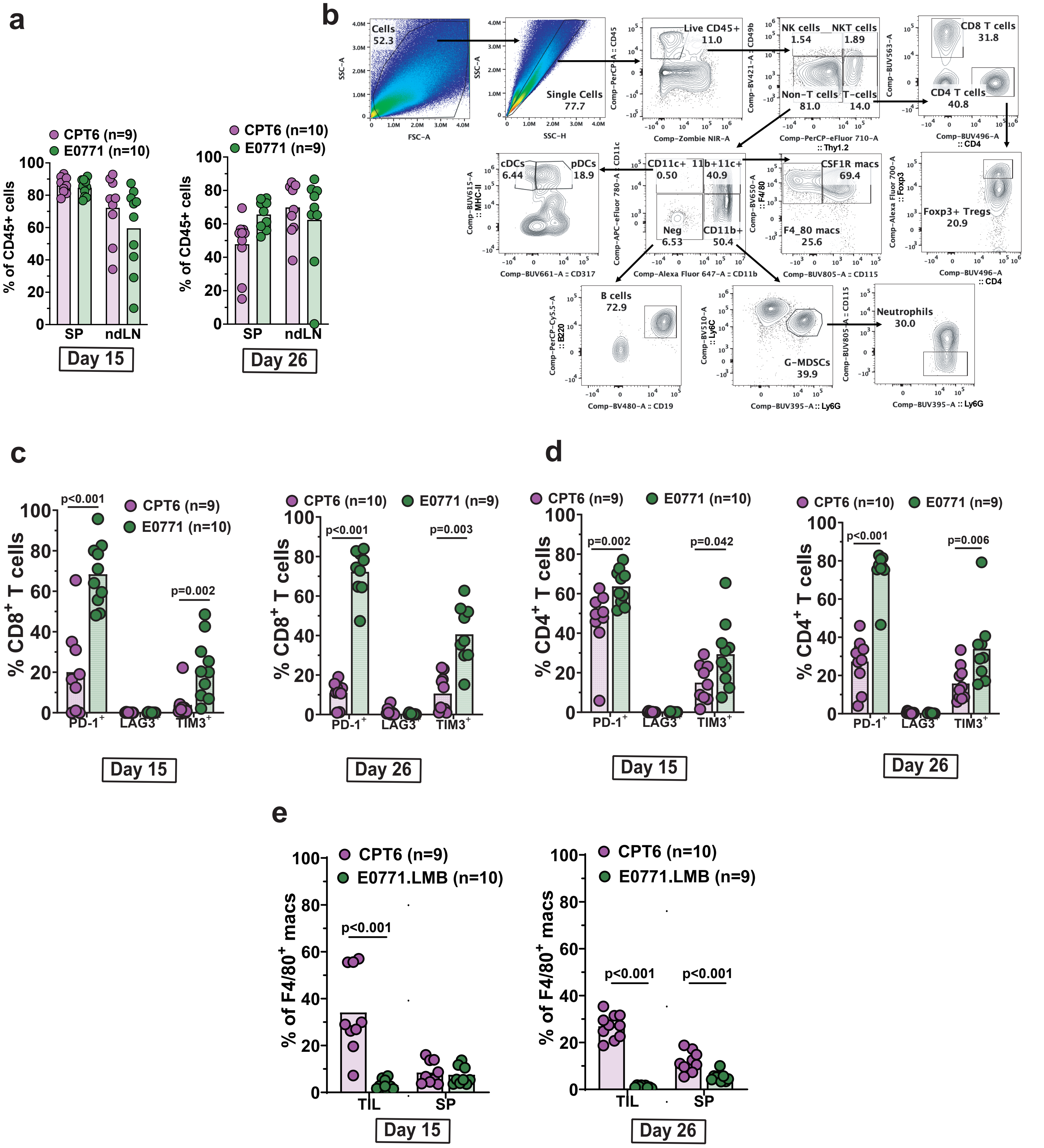


**Figure S4. Immune infiltration landscape of tumor microenvironment in CPT6 and E0771 *in vivo* tumor. a.** Percentage of CD45+ cells in spleen and lymph node after CPT6 and E0771 cell injection. **b.** Flow cytometry workflow in identifying immune compositions in immune microenvironment analysis. **c.** Bar graphs depicting percentage of tumor infiltrating CD8 T cells expressing PD-1, LAG3 and TIM3 at D15 (left panel) and D26 (right panel) comparing CPT6 and E0771. **d.** Bar graphs depicting percentage of tumor infiltrating CD4 T cells expressing PD-1, LAG3 and TIM3 at D15 (left panel) and D26 (right panel) comparing CPT6 and E0771. **e.** Proportion of CX3CR1+ F4/80+ macrophages in tumor compared to spleen in CPT6 and E0771 at D15 (left panel) and D26 (right panel).
